# Modeling ecological communities when composition is manipulated experimentally

**DOI:** 10.1101/2022.05.12.491213

**Authors:** Abigail Skwara, Paula Lemos-Costa, Zachary R. Miller, Stefano Allesina

**Affiliations:** Department of Ecology & Evolution, University of Chicago, 1101 E. 57th st., Chicago, IL 60637, USA; Department of Ecology & Evolutionary Biology, Yale University, New Haven, CT; Nortwhestern Institute for Complex Systems, Northwestern University, Evanston, IL

**Author notes:** Correspondence: Abigail Skwara < >.

**Keywords:** Generalized Lotka-Volterra model, Generalized Linear model, weighted least squares, species interactions, species coexistence

## Abstract

1. In an experimental setting, the composition of ecological communities can be manipulated directly. Starting from a pool of *n* species, one can co-culture species in different combinations, spanning mono-cultures, pairs of species, and all the way up to the full pool. Here we advance methods aimed at inferring species interactions from data sets reporting the density attained by species in a variety of sub-communities formed from the same pool.
2. First, we introduce a fast and robust algorithm to estimate parameters for simple statistical models describing these data, which can be combined with likelihood maximization approaches. Second, we derive from consumer-resource dynamics statistical models with few parameters, which can be applied to study systems where only a small fraction of the potential sub-communities have been observed. Third, we show how a Weighted Least Squares (WLS) framework can be used to account for the fact that species abundances often display a strong relationship between means and variances.
3. To illustrate our approach, we analyze data sets spanning plants, bacteria, phytoplankton, as well as simulations, recovering a good fit to the data and demonstrating the ability to predict experiments out-of-sample.
4. We greatly extend the applicability of recently proposed methods, opening the door for the analysis of larger pools of species.

## Introduction

Efforts to relate empirical measurements of population growth to dynamical models can be traced back to the origins of ecology. For example, in his famous study proposing the logistic growth model, Verhulst (1838) presented the fitted time-series for the human population growth of France and Belgium. Interest in contrasting empirical data with models for population dynamics has only grown through the years (Gilpin, 1973; Powell & Steele, 2012), with the development of sophisticated approaches (Ellner et al., 2002) accounting for different sources of errors (Carpenter et al., 1994; De Valpine & Hastings, 2002), and important applications such as forecasting population trajectories (Clark et al., 2001; Sugihara et al., 2012), modeling the evolution of disease epidemics (Du et al., 2017), and detecting chaos in natural systems (Perry et al., 2012; Sugihara & May, 1990).

Because of the intrinsic difficulty of manipulating natural systems, much of the literature on these issues has historically focused on inferring the parameters of dynamical models from time-series observations (Downing et al., 2020; Ives et al., 2003; Vandermeer, 1969). With the advent of high-throughput laboratory techniques, however, a different approach has become viable. Instead of attempting to learn the parameters of a model by analyzing the fluctuations of several interacting populations through time, it is instead possible to use steady-state abundances recorded for a variety of community compositions, each captured at a single time point, to infer model parameters (Ansari et al., 2021; Fort, 2018; Maynard et al., 2020; Voit et al., 2021; Xiao et al., 2017). For a species pool of interest, the initial species composition is manipulated in a series of experiments, and then the resulting set of final community compositions can be used to estimate the parameters of a statistical model. For example, to infer the interactions among *n* species, one could perform a series of experiments in which either a single species is grown in isolation, or in which pairs, triplets, or larger subsets of species are co-cultured (Dormann & Roxburgh, 2005; Friedman et al., 2017). Once any transient dynamics have elapsed, densities of all surviving species are recorded. Provided that a sufficiently large and diverse set of sub-systems (i.e., distinct species compositions) have been observed, it is possible to infer the parameters of a statistical model for species interactions from these static measurements Maynard et al. (2020). Such statistical models can be derived from corresponding dynamical models, thereby linking static estimates to models for population dynamics.

We build upon previous work (Carrara et al., 2015; Maynard et al., 2020; Xiao et al., 2017) in which the density of each species is assumed to be linearly related to the densities of the other species in a community. This statistical structure arises naturally from the ubiquitous Generalized Lotka-Volterra (GLV) dynamical model (Lotka, 1920; Volterra, 1926)—although it is not necessary for dynamics to obey this simple model to yield a good fit to data (Maynard et al., 2020). While very easy to formulate, this type of statistical model is both difficult and computationally expensive to fit. Here, we extend the approach of Maynard et al. (2020) in three main ways to overcome issues that have limited the application of these methods. First, we introduce a fast iterative algorithm that finds parameters yielding a good fit to data. This algorithm can be used in conjunction with more rigorous — but less efficient — fitting approaches by providing a high-quality starting point for numerical likelihood optimization. We show that this hybrid approach is computationally very efficient and produces better results than current methods. Second, we derive simplified versions of the statistical model by exploiting the parallel between the model structure and Lotka-Volterra dynamics.

These versions of the model require the estimation of fewer parameters, while still providing a good fit to data and retaining a clear ecological interpretation. Finally, we show that a more sophisticated error model for the observations results in a Weighted Least Squares problem that can be solved efficiently. Extending the error structure offers more flexibility to fit empirical data, especially when the variance of experimental observations changes with the mean, as typically seen in ecological data (Grilli, 2020; Taylor, 1961). To illustrate these improvements, we examine four recently published data sets spanning communities of plants (Kuebbing et al., 2015), phytoplankton (Ghedini et al., 2022), and bacteria (Chu et al., 2021), as well as simulated data.

## Materials and Methods

### Experimental setup and data

Given a pool of *n* species, suppose we have performed a large set of experiments in which different combinations of species are co-cultured for a suitably long time. At the end of each experiment (once transient dynamics have elapsed), we measure and record the biomass/density of each extant species. The experiments encompass a variety of initial compositions and initial densities, and are conducted with replication. Provided that our measurements, after any species extinctions, span a sufficient variety of sub-communities, we can fit a simple statistical model to these empirical “endpoints,” which may in turn be used to predict the densities of each species in any unobserved subset, and in particular whether the given subset of species will coexist.

To test our models, we use four recently-published data sets that have a suitable experimental design. In particular, we consider two data sets from Kuebbing et al. (2015), who selected two pools (natives, non-natives) of four plants each, and grew them in 14 out of 15 possible combinations of species. Similarly, Ghedini et al. (2022) considered five phytoplankton species, and grew them in monoculture, in all possible pairs, and all together. Finally, we consider a subset of the data from Chu et al. (2021), consisting of communities of four bacterial strains co-cultured along with *Pseudomonas fluorescens* in different combinations. A detailed description of each data set is reported in the Supporting Information S1.

### A simple statistical framework

We start by summarizing the statistical framework presented in Maynard et al. (2020), which we will extend below. The framework assumes that we have measured densities for several of the possible communities of coexisting species we can form from a pool of *n* species. The approach rests on two main assumptions. First, we take each observed measurement to be a noisy realization of a “true” value:

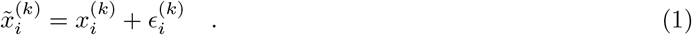

That is, the observed density of species *i* when grown in community *k*, 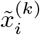, is given by a “true” value, 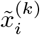, modified by an error term, 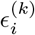. We can arrange our empirical data in a matrix 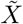, with *n* columns (one for each species in the pool) and as many rows as the number of observed communities. In particular, 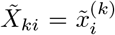 if species *i* is present in community *k*, and zero otherwise. Similarly 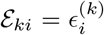 if species *i* is in community *k* and zero otherwise. Then, we can rewrite Eq. 1 in matrix form:

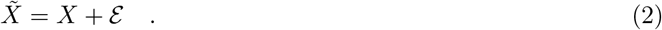

When several replicates are available, we assume that the density recorded for species *i* in community *k* and replicate *r* follows 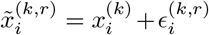, i.e., all replicate measurements of species *i* in community *k* stem from the same true value (to recover the notation in Eq. 2, we simply stack the replicates in matrix 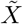 and the corresponding true means in matrix *X*). This assumption amounts to ruling out “true multistability” in the underlying dynamics (Xiao et al., 2017) — if a set of species coexists, it must always reach the same attractor (be it an equilibrium, cycle, or chaotic attractor). Notice, however, that this framework is compatible with the fact that experiments initialized with the same set of species may yield distinct coexisting communities. This could occur if stochastic dynamics or differences in initial densities drive two experimental systems to different attractors. Because this framework exclusively uses data gathered at the end o f e ach experiment, it is completely blind to initial conditions; we only require that communities that reach the same final composition have also reached the same dynamical attractor.

Second, following Maynard et al. (2020), we assume the true species’ densities in a given community are linearly related to each other. For any species *i* in community *k*, we can express these densities by writing:

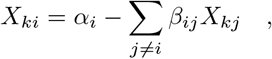

where *α_i_* is the density that species *i* would attain when grown in isolation, and the coefficients *β_ij_* model the effects of the other species in *k* on the density of species *i*. Importantly, *β_ij_* depends only on the identities of the species, and not on the community we are modeling—this assumption amounts to ruling out higher-order interactions or other nonlinearities that would make the per-capita effect of species *j* on species *i* dependent on the state of the system. Rearranging, we obtain:

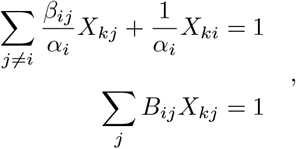

with *B_ii_* = 1/*α_i_* and *B_ij_* = *β_ij_/α_i_*. Naturally, systems with sufficiently strong higher-order effects (or highly nonlinear systems) would be incompatible with this assumption. Indeed, Maynard et al. (2020) used simulations to show that, while the framework is generally quite robust to model misspecification, strong higher-order interactions resulted in poor fit and poor inference of true parameters. Conversely, good fit to the data would indicate the lack of strong higher-order interactions or nonlinearities.

Because *X_kj_* = 0 whenever species *j* is not present in community *k*, we can define the sub-matrix *B*^(*k*)^ obtained by retaining only the rows and columns of *B* corresponding to species that are present in community *k*. We similarly take *X*^(*k*)^ to be a vector containing only the densities of the species in *k*, and 1^(*k*)^to be a vector of ones with as many elements as *X*^(*k*)^. Then the model can be written in matrix form as *B*^(*k*)^*X*^(*k*)^ = 1^(*k*)^for each community *k*.

Suppose that we have obtained a matrix *B* and we want to make a prediction about a set of species *k*, such as whether these species can coexist. We solve the corresponding equation for *X*^(*k*)^,

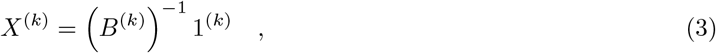

with two possible outcomes: a) all the components in *X*^(*k*)^ are positive, in which case we conclude that the species can coexist, with predicted densities *X*^(*k*)^; or b) some of the components in *X*^(*k*)^ are negative, which we interpret as the impossibility of coexistence of this combination of species (Maynard et al., 2020).

Arguably, the simplest version of this statistical model is obtained by assuming that errors are independent, identically-distributed random values sampled from a normal distribution, such that 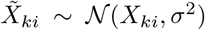 whenever species *i* is in community *k*. Then, fitting the model requires minimizing the sum of squared deviations between the observed data and model predictions (Ordinary Least Squares, OLS).

This suggests a straightforward method to fit the model: propose a matrix *B*, compute the predicted densities for all species in all observed communities using Eq. 3, and search for the matrix *B* that minimizes the deviations between the observed data 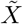 and the predicted *X*. Unfortunately, this method is quite inefficient and computationally very expensive, as it requires inverting a sub-matrix of *B* for each observed community, and there may be up to 2^*n*^ − 1 unique community compositions that can be built from a pool of *n* species. Moreover, the problem of minimizing deviations is markedly non-convex—starting from different initial estimates, we are likely to converge to different (and thus in general sub-optimal) solutions.

To circumvent this problem, Maynard et al. (2020) proposed a simple analytical approach (dubbed the “naïve method”) to find a rough estimate of *B* from observed data; this initial estimate of *B* can then be used as a starting point for more sophisticated fitting routines. However, as discussed in their study, this method suffers from a number of issues (detailed in Supporting Information, S3). In particular, the statistical model assumes that the observations are noisy, while the naïve method assumes that they have been observed precisely and that instead Eq. 3 only holds approximately. In practice this means that this approach does not provide the maximum likelihood estimate for *B* when the data are noisy. One of the goals of the present work is to build an iterative algorithm that is not only computationally efficient, but that also improves upon the naïve estimate for *B* by yielding a superior starting point for subsequent parameter search.

## Results

### A fast, iterative algorithm to estimate parameters

We want to find the maximum likelihood estimate for the matrix *B*, i.e., the choice of *B* minimizing the sum of squared deviations:

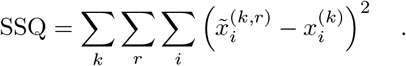

The computational bottleneck we face is that determining the predicted abundances 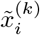 from the matrix *B* (via Eq. 3) for all community compositions is very expensive (requiring the inversion of many matrices). We therefore develop an algorithm in which this expensive calculation is performed only seldomly and at defined intervals, and optimization is carried out in between these steps without having to re-calculate the predicted 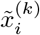. To achieve this goal, we divide the process of optimizing *B* into two steps: a prediction step (computationally expensive), and a numerical optimization step (computationally cheaper). By alternating between the two steps, we quickly converge to a good draft matrix *B*.

To construct this algorithm, we derive two “auxiliary” matrices that are useful for computation. Having arranged our data in the matrix 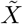 as detailed above, we take 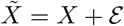 (Eq. 2), transpose each side, and multiply by *B*. We obtain the sum of two new matrices:

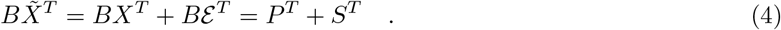

In the remainder of this section we use *P* and *S* simply as a convenient means to build our algorithm—we discuss their ecological interpretation in Supporting Information S4. From Eq. 4, we have *B*^−1^*P^T^* = *X^T^* and *B*^−1^*S^T^* = E ^*T*^. Our sum of squared deviations is simply the squared Frobenius norm of ε, 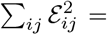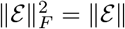, and from Eq. 2 and 4, we can write:

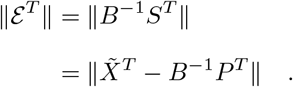

Our goal is therefore to find a matrix *B* such that *B*^−1^*P^T^* is as close as possible to the observed data 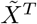. However, neither *B* nor *P* are known, although *P* can be calculated from *X* and *B*, and hence from *B*. We therefore attempt to minimize the sum of squared deviations through an iterative algorithm reminiscent of the Expectation-Maximization (EM) approach (Moon, 1996):

1. Propose a candidate matrix *B*;
2. Consider *B* to be fixed, and use it to compute *X* via Eq. 3 (prediction step);
3. Compute *P^T^* = *BX^T^*;
4. Consider *P* to be fixed, and find an updated *B*^−1^ by numerically minimizing 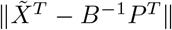 (optimization step). Invert it to recover an updated *B*;
5. Repeat steps 2-4 until convergence.

In principle, this iterative algorithm depends on the proposal of the initial matrix: for our numerical experiments, we always start from the identity matrix (i.e., no interactions between species). Finally, we perform an additional numerical optimization to refine the results produced by the algorithm. The algorithm above may return values of *X_ki_* < 0, because the solution minimizing the sum of squared deviations does not need to contain only non-negative densities. Clearly, such solutions would be biologically unattainable, and qualitatively incompatible with the observed data. Thus, when numerically refining the solution, we only accept proposals that yield non-negative predictions for the observed densities.

While this algorithm is not guaranteed to find the maximum likelihood estimate of *B*, we observe that in practice, the combination of our iterative algorithm and the final numerical optimization step yields very good solutions. In all cases, we converge on a solution that is superior to the result using the naïve method. As shown in Figure 1 (and Supporting Information, S8), the SSQ typically decays rapidly with the number of iterative steps, and the final numerical search provides an additional improvement. While for the data presented here the algorithm converges smoothly, in principle, one could observe the SSQ oscillating as the algorithm progresses. As for gradient descent and similar methods, this problem can be alleviated by introducing a “momentum” (Polyak, 1964; Qian, 1999), i.e., when updating matrix *B* in Step 4, one could take a weighted average of the matrix used to compute *P* (the current estimate of *B*), and the matrix that maximizes the likelihood given *P* (the new estimate of *B*). The addition of momentum would help achieve a smooth, monotonic decay of the SSQ, at the cost of having to take a larger number of steps before convergence.

**Figure 1:**
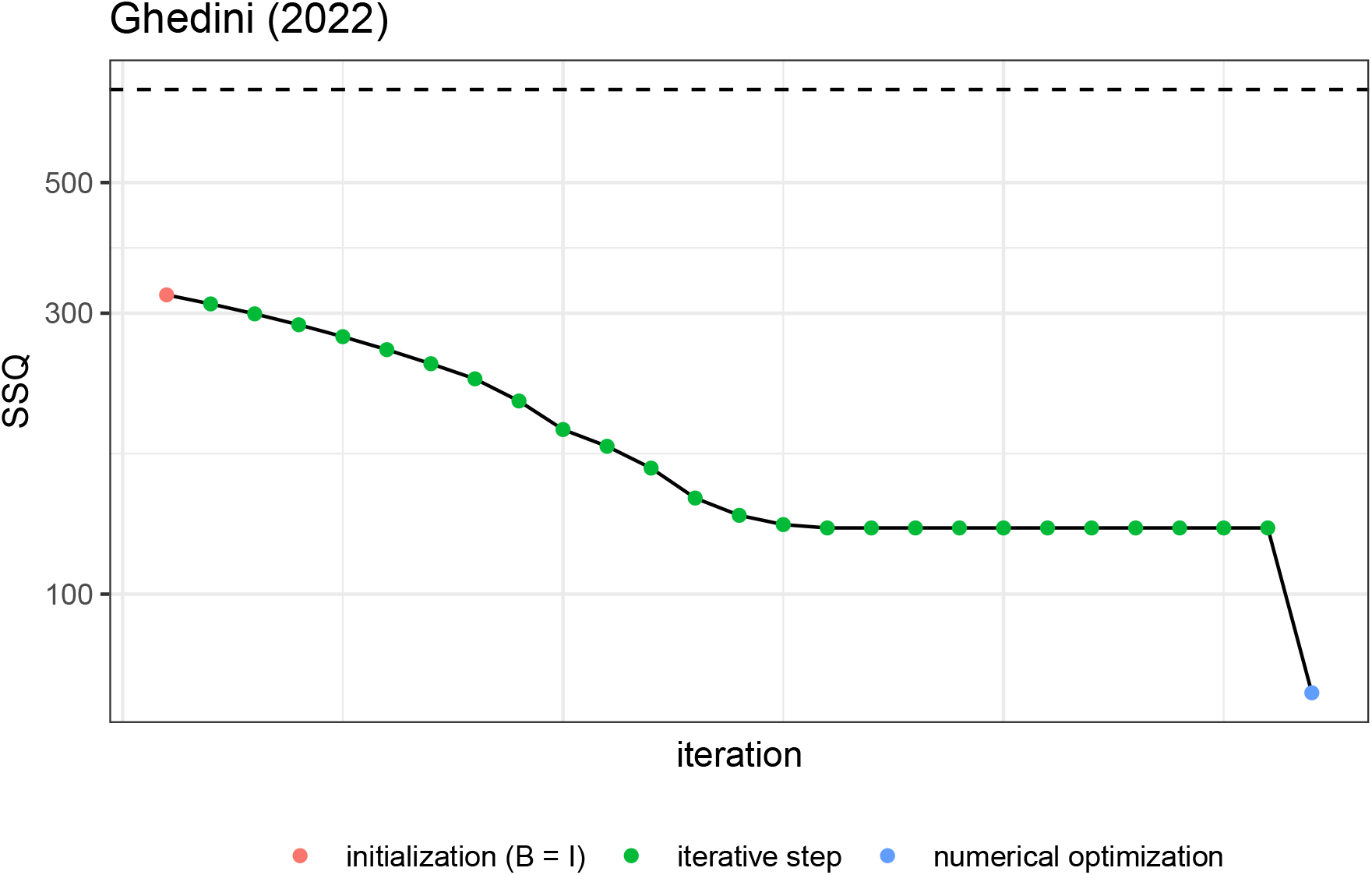
Sum of squared deviations (shown in log-scale, y-axis) for the initial matrix *B* = *I*, at each successive iteration of the algorithm, and after the numerical maximization (colors). The data from Ghedini et al., 2020. is used as an example. The dashed line marks the SSQ obtained using the naïve method of Maynard et al., 2020.

### Simplified versions of the model

Fitting the *n*^2^ parameters of the interaction matrix *B* requires having observed a sufficiently varied set of experimental community compositions. It is necessary to observe each of the *n* species in at least *n* distinct communities, and each pair of species coexisting in at least one community (for a full derivation of these conditions, see Supporting Information, S2). These are very demanding requirements, and therefore many published data sets do not contain a sufficient variety of communities to allow the identification of all parameters. These conditions grow more onerous with the number of species in the pool, making the approach unfeasible for species pools of even moderate size. To address this issue, we propose a nested set of simplified versions of the statistical model that use fewer parameters, but retain the basic model structure and a straightforward ecological interpretation. The data requirements are greatly reduced, scaling linearly, rather than quadratically with *n* (see Supporting Information S7 for detailed data requirements). These simpler models have the added benefit of being extremely efficient to fit from a computational standpoint (Supporting Information, S6), and useful for model selection, especially when one suspects that the full model with *n*^2^ parameters may be susceptible to overfitting.

We follow a disciplined approach to develop these simpler models: we consider a version of the MacArthur’s consumer-resource model (MacArthur, 1970) in which each species has access to its own private resources, and all species have access to a shared resource (Supporting Information, S5). By iteratively reducing the number of free parameters in the model, we obtain simpler structures for the matrix *B* (Table 1).

**Table 1:**
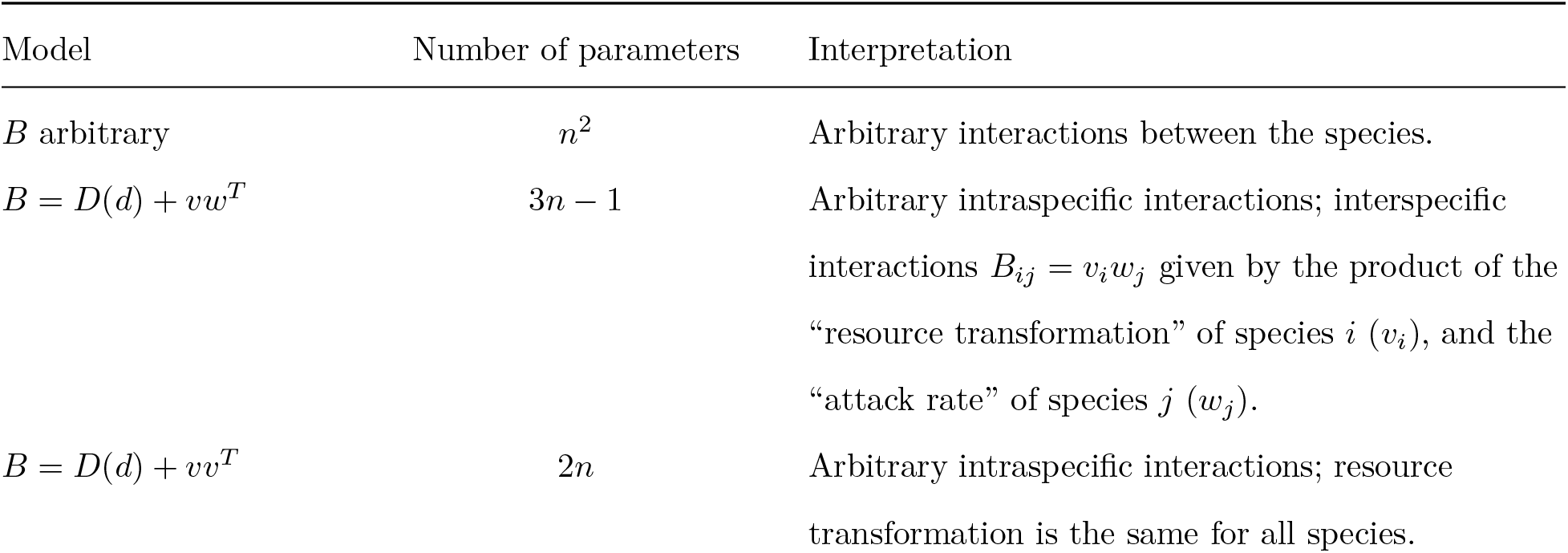
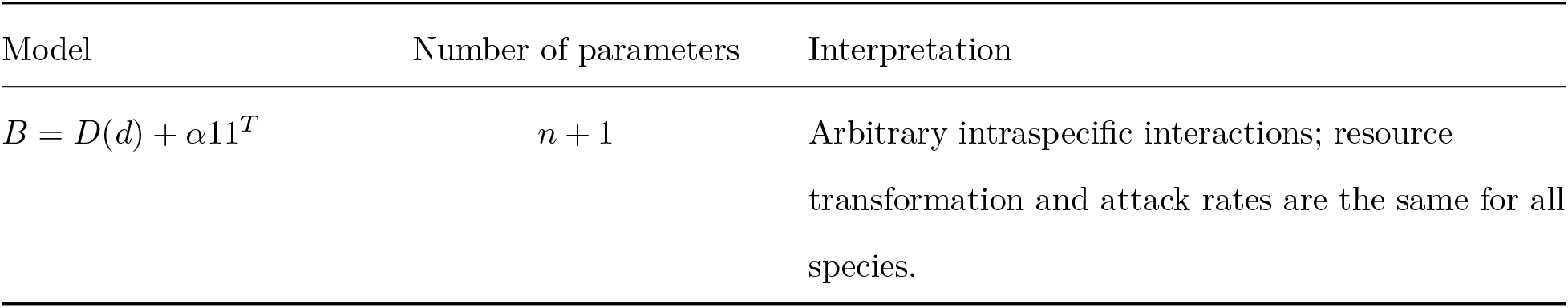
Simpler versions of the model derived by considering a MacArthur’s consumer-resource model in which species have access to a private pool of resources as well as a shared pool of resources. See Supporting Information, S5 for a derivation.

| Model | Number of parameters | Interpretation |
| --- | --- | --- |
| $B$ arbitrary | $n^2$ | Arbitrary interactions between the species. |
| $B = D(d) + vw^T$ | $3n - 1$ | Arbitrary intraspecific interactions; interspecific interactions $B_{ij} = v_i w_j$ given by the product of the “resource transformation” of species $i$ ($v_i$), and the “attack rate” of species $j$ ($w_j$). |
| $B = D(d) + vv^T$ | $2n$ | Arbitrary intraspecific interactions; resource transformation is the same for all species. |
| $B = D(d) + \alpha 11^T$ | $n + 1$ | Arbitrary intraspecific interactions; resource transformation and attack rates are the same for all species. |

Given that the matrix *B* can be interpreted as detailing pairwise interactions (i.e., the effect of the density of species *j* on species *i*), the model *B* = *D*(*d*) + *vw^T^* corresponds to the following ecological picture: each species interacts with conspecifics through their private resources (corresponding to the coefficients *d_i_*) and with all species via the shared resource. Interactions arising from the shared resource are given by the product of a resource utilization vector *w* (i.e., attack rates), and a resource transformation vector *v*, where each species is characterized by its *v_i_* and *w_i_* values. The interpretation of the simpler models is similar: by considering equal transformation rates one makes *vw^T^* = *vv^T^* symmetric, and by assuming that all species also have the same attack rates, *vw^T^* = *α*11^*T*^. Note that in all these simplified models, the intraspecific interactions (i.e., the diagonal elements of *B*) are modeled with great flexibility; the reduction in parameters is obtained by assuming that interspecific interactions follow a simple pattern, defined by few species’ traits.

In Figure 2, we show the fit of these four models when analyzing the data reported in Ghedini et al. (2022).

**Figure 2:**
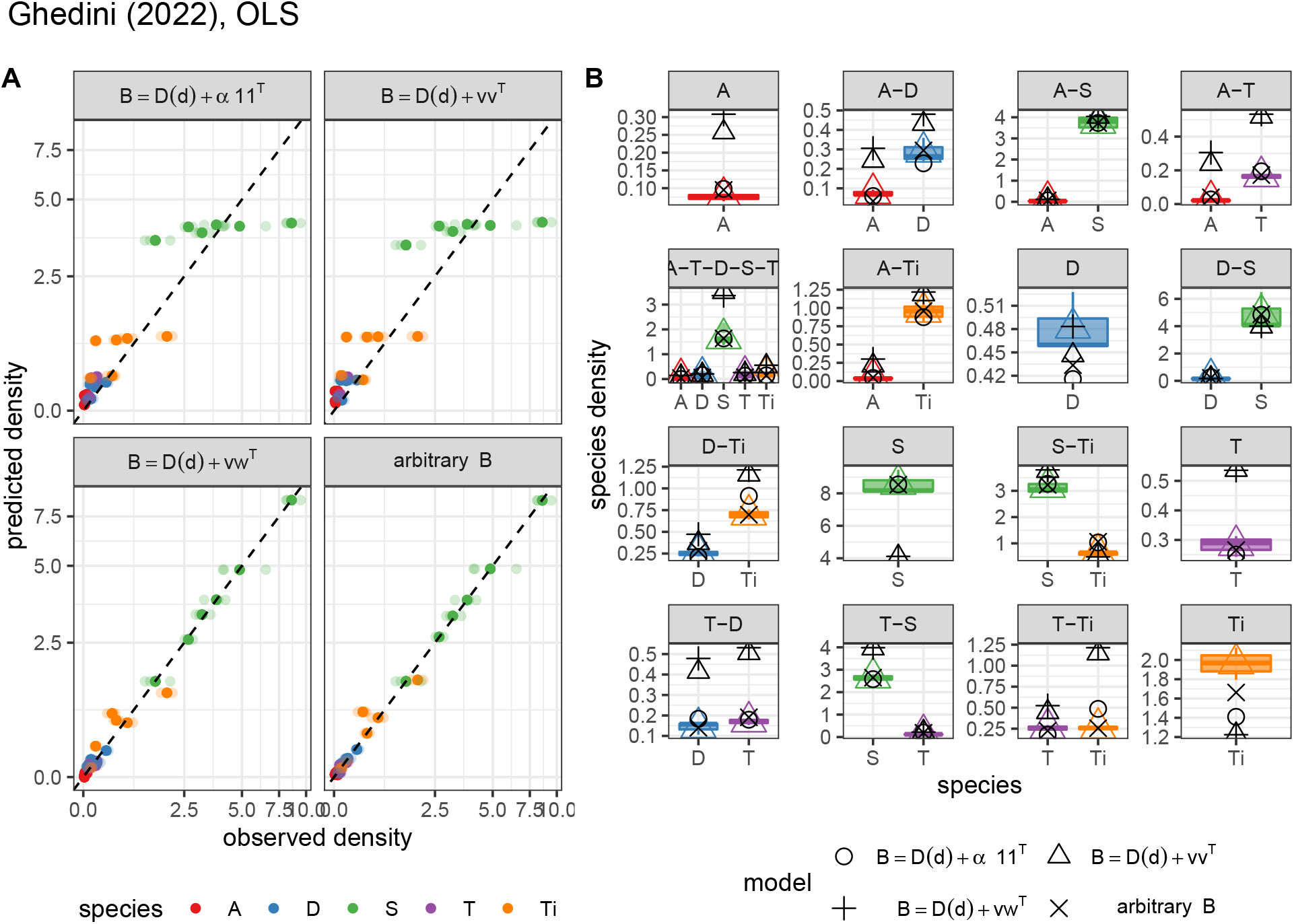
A. Observed species (color) densities (x-axis) vs. predicted densities (y-axis) for the data by Ghedini et al. (2022), when fitting the four version of the model (panels). Replicate measurements for each species/community are reported as semi-transparent points; the mean for each species/community combination as a solid point. B. Observed species (color, x-axis) densities (y-axis). Boxplots show the distribution of the species densities across replicates, with the median density reported as a solid colored line; the mean density is represented by the colored triangle. Predicted values for each species in each community are represented by black open symbols (one for each of the four versions of the model).

For all data sets we find the same qualitative results: the full model (*n*^2^ parameters, 25 parameters for the data set of Ghedini et al. (2022)) and the simplified model in which only two values per species determine all interspecific interactions (3*n* − 1 parameters, 14 for Ghedini et al. (2022)) have very similar performance (Supporting Information, S8), while any further simplification results in a marked loss of fit. These trends are evident when contrasting the total SSQ across models and data sets (Figure 3).

**Figure 3:**
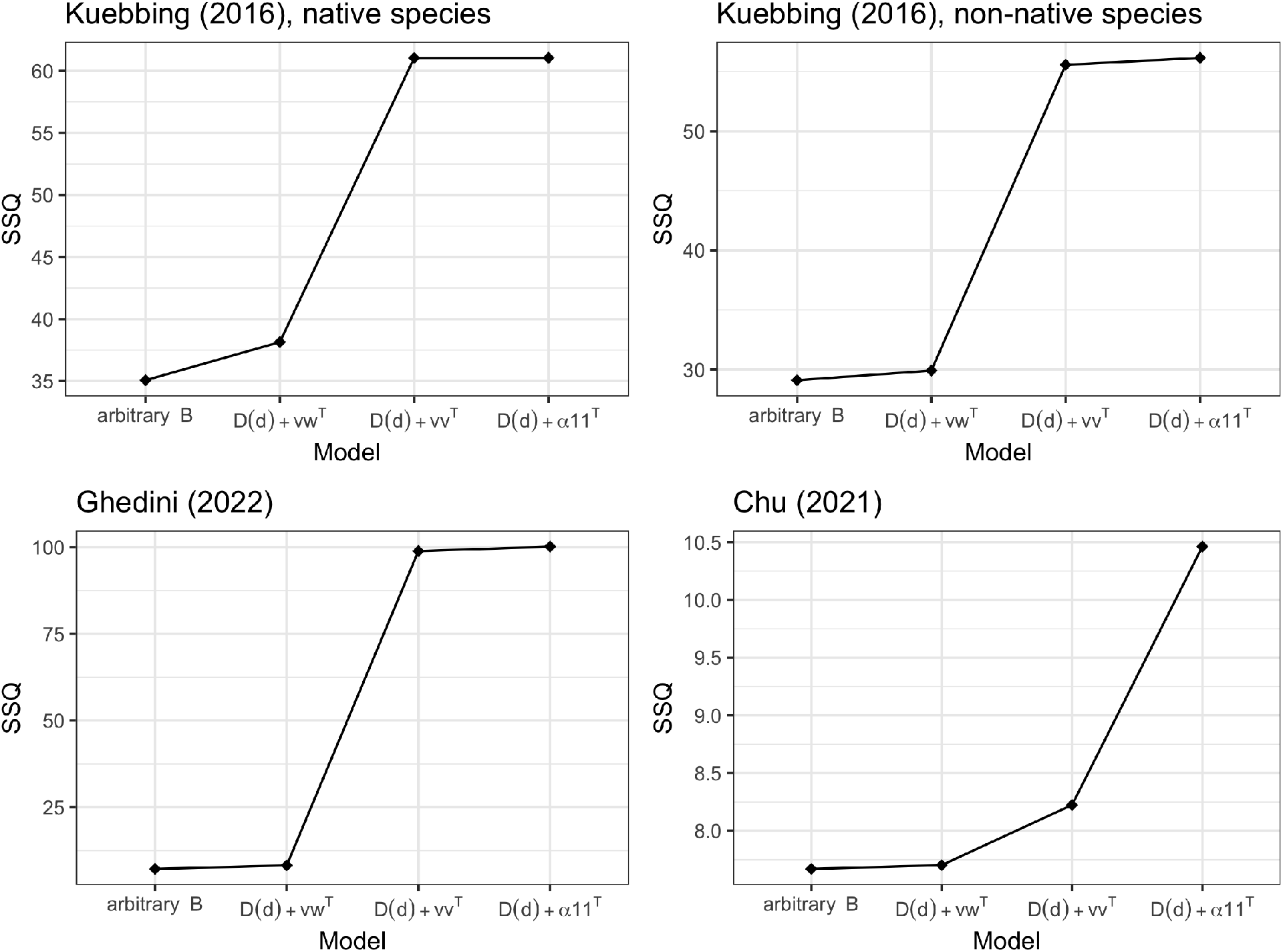
Sum of squared deviations between observed and predicted densities for the four data sets considered, fitted using the four models in Table 1.

### Allowing the variance to change with the mean

So far, we have assumed that errors are independent and identically-distributed for all measurements. In many situations, however, this assumption would be quite unrealistic. For example, some species could systematically grow to much higher density than others—resulting in potentially larger absolute errors in their measurement—or measurements might be made on a small sub-plot or sample volume and then extrapolated to the whole plot or volume through multiplication. In such cases, the data would display marked heteroskedasticity (i.e., the variance would change with species density).

In Figure 4, we show that the variance scales with a power of the mean in the phytoplankton data from Ghedini et al. (2022). This is in fact the expected behavior of many ecological systems, as posited by Taylor’s law (Gaston & McArdle, 1994; Routledge & Swartz, 1991; Taylor, 1961).

**Figure 4:**
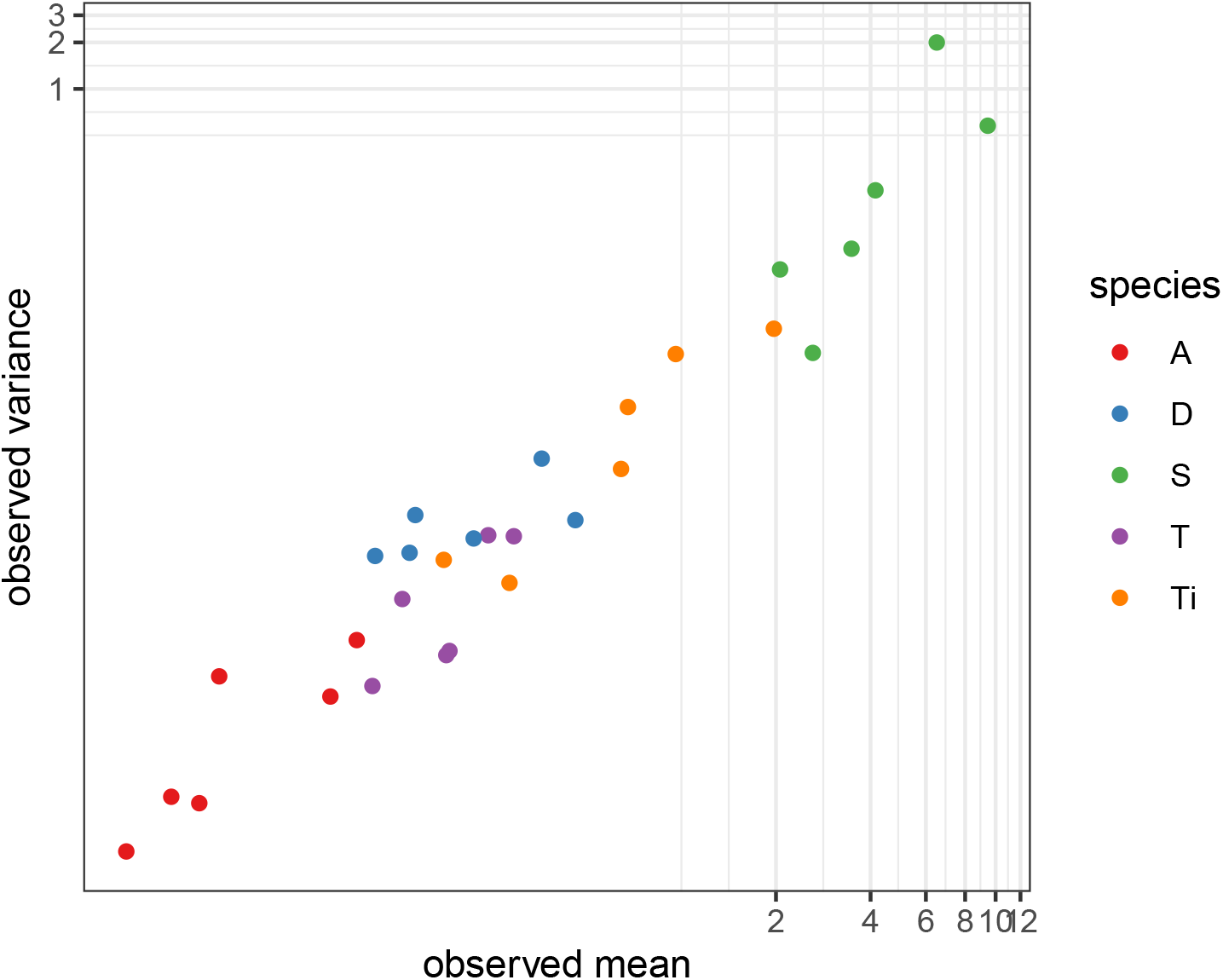
Mean (x-axis) vs. variance (y-axis) for the data published by Ghedini et al., 2022. For each species (colors) and community combination, the mean and variance of the observed species densities are computed across replicates. Note that both axes have a logarithmic scale, and thus the strong linear trend displayed by the data corresponds to a power-law relationship between the mean and the variance.

A possible approach to dealing with the systematic heteroskedasticity observed in these data sets is to use Weighted Least Squares (WLS) instead of OLS. To implement this approach, each squared residual is divided by the variance of the corresponding distribution, which is equivalent to measuring residuals as standard deviations of the standardized data. Here for simplicity we use the variance in the observed 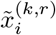 to estimate the variance 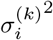 (a more sophisticated approach would call for the concomitant estimation of means and variances, as in methods based on iteratively reweighted least squares). The WLS approach can be interpreted as reweighting the relative importance of errors made in our predictions, such that small errors made when estimating low species abundances are penalized as much as larger errors made when predicting higher species abundances. This is in contrast to OLS, where a one percent error in estimating the density of a species with high abundance contributes much more to the sum of squared deviations than the same proportional error for a species with low abundance.

Since experimental replicates are necessary to estimate these variances, we must have replicates for all communities (as we do for all the data sets considered here) in order to implement the WLS approach in this straightforward manner.

In Figure 5 (and Supporting Information, S8), we compare these two error structures and find that we indeed observe a better fit for species with lower abundances when the WLS approach is used. This has important implications for the qualitative prediction of species’ coexistence, because for a species present at low abundance in a community, the numerical difference between a positive abundance and a negative abundance (equivalent to a prediction of lack of coexistence) could be quite small.

**Figure 5:**
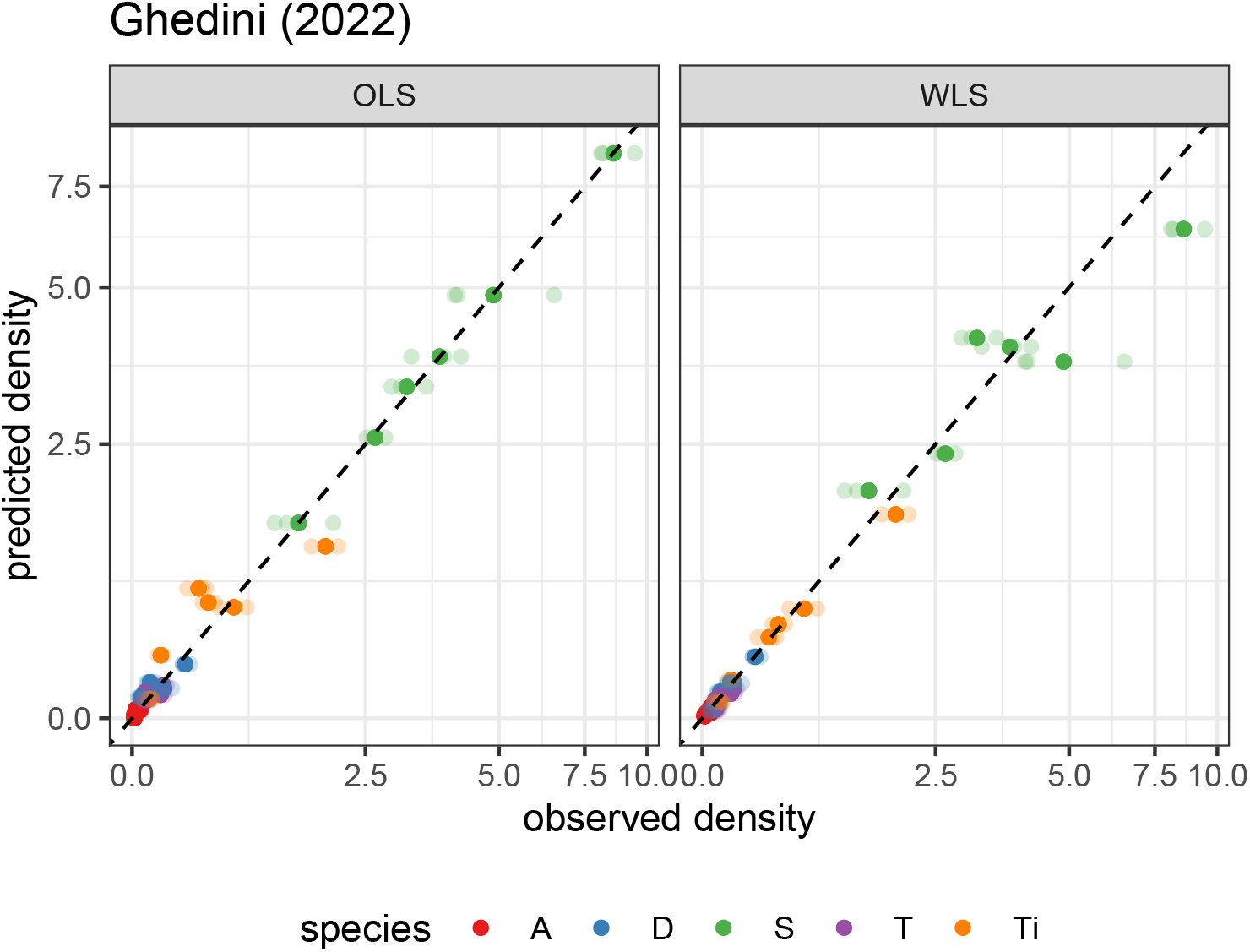
Observed (x-axis) vs. predicted (y-axis) species densities in all communities using the data by Ghedini et al. (2022), when fitting the simplified model *B* = *D*(*d*) + *vw^T^* and either minimizing the sum of squared deviations (OLS) or the sum of standardized squared deviations (WLS). Replicate measurements for each species/community are reported as semi-transparent points; the mean for each species/community combination is reported as a solid point. In OLS, small (proportional) deviations of highly abundant species (e.g., *Synechococcus* sp., in green) are penalized more than larger (proportional) deviations of lower-abundance species (e.g., *Tisochrysis lutea*, in yellow). In contrast, when performing WLS each data point is standardized by its corresponding variance, leveling the importance of each measurement.

Finally, in Supporting Information S9 we perform simulations in which observations are taken from distributions with known mean-variance relationship, and fit the data using OLS, WLS or a likelihood-based approach. In all cases, we find that WLS outperforms OLS, allowing us to fit the data well and recover with good confidence the parameters used to generate the data.

### Predicting experiments out-of-sample

For the empirical data sets considered here, we always find good agreement between the observed and the fitted values when using the full model under either an OLS or WLS scheme. We also consider a leave-one-out (LOO) cross-validation approach, to verify that the models capture real features of the interactions between species in the communities, and are not simply over-fitting the data. For a data set in which *m* experimental communities have been measured, we implement the LOO approach by designating one of the communities (along with any replicates) as out-of-sample (sometimes called “out-of-fit,” as in Maynard et al. (2020)), and fitting our model on the remaining *m* − 1 communities. This process can be repeated for each of the *m* communities in turn, and we then compare the predicted species’ abundances with their experimentally observed values, as shown in Figure 6 (and Supporting Information, S8). While the quality of the predictions is necessarily worse, in almost all cases we would have correctly predicted the experimental outcome both qualitatively (i.e., possibility of coexistence) and quantitatively.

**Figure 6:**
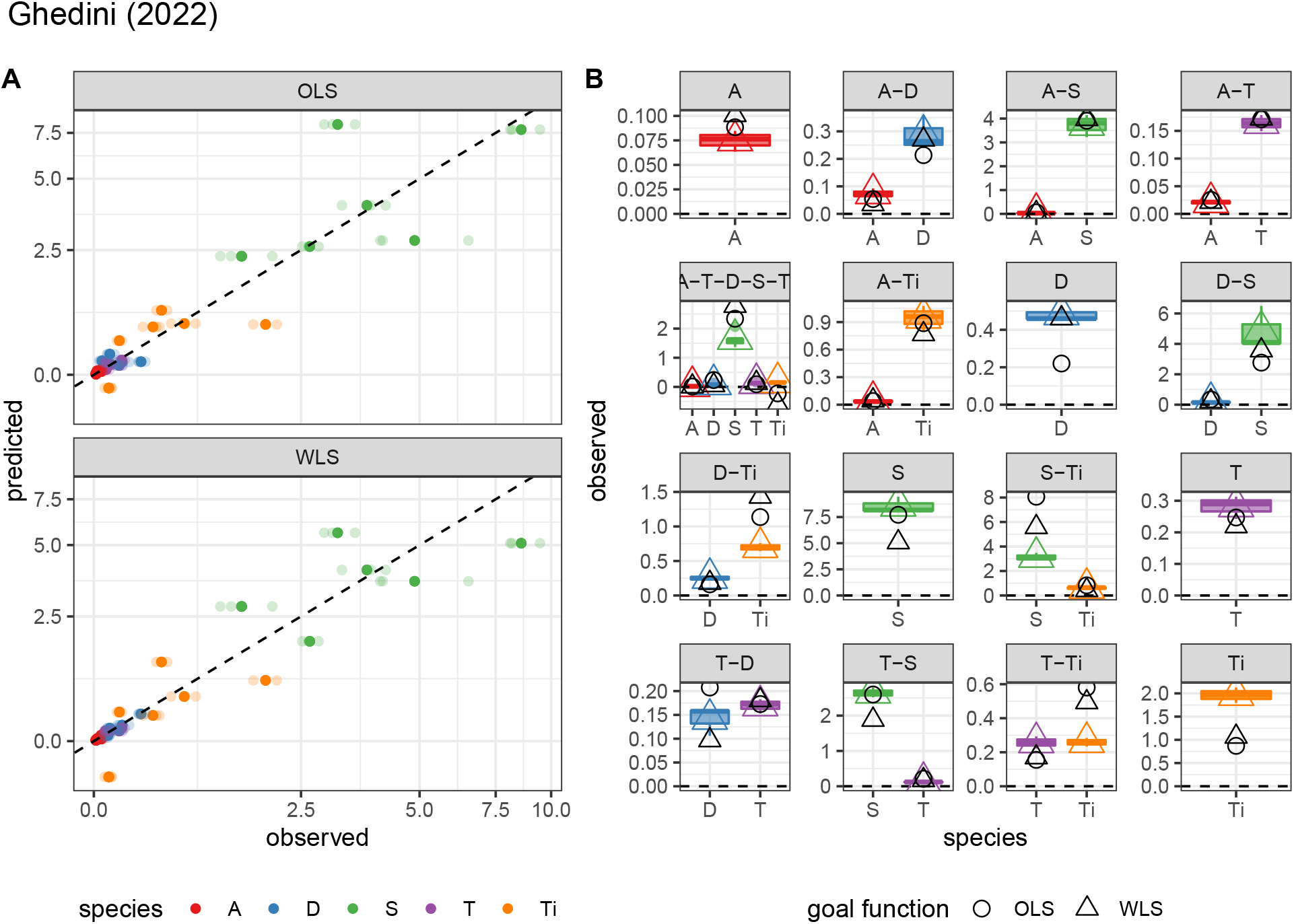
As Figure 2, but showing only out-of-sample predictions. Each panel is obtained by excluding the corresponding data, fitting the model using the remaining data, and then predicting the data reported in the panel out-of-sample. Results are presented for the model *B* = *D*(*d*) + *vw^T^* under either OLS or WLS. Note that in the panel displaying the measurements for the full community (A-T-D-S-Ti), *Tisochrysis lutea* is predicted at a negative abundance—without having observed these data, our models would suggest a lack of coexistence.

## Discussion

In this study, we have further developed a simple, extensible framework to infer species’ interactions from experimental data where community composition has been manipulated, and showcased its potential by testing this approach on data sets spanning plant, phytoplankton, and bacterial communities, as well as simulations. We have improved the computational performance of the fitting routines, derived simplified versions of the model, and extended the error structure used to capture the variability in the data.

Previous studies have used a variety of approaches to model this type of experimental data, including bottom-up “assembly rules,” in which the focus is on observing all possible pairs of species and using the outcomes to predict coexistence of larger species assemblages (Dormann & Roxburgh, 2005; Friedman et al., 2017); methods relying on monocultures and leave-one-out communities (Ansari et al., 2021; Venturelli et al., 2018), using the extremes of very simple and highly speciose communities to infer pairwise interactions; and methods based on time-series that use fluctuations to estimate interaction strengths and the effects of environmental variables on ecological dynamics (Downing et al., 2020; Ives et al., 2003).

Our framework bridges the dynamical and statistical perspectives of these approaches. Importantly, though the models presented here are statistical in nature, they have a straightforward connection to the Generalized Lotka-Volterra dynamical model. By exploiting this parallel, we have derived a family of simplified models that put a premium on the ecological interpretability of the parameters. While an alternative approach to model regularization would be data-driven, machine learning methods (e.g., enforcing the sparsity or parsimony of *B* through a penalized regression), we have shown that ecologically-motivated model constraints are a viable option. By clearly relating statistical and dynamical models, we retain the ability to probe other properties of these systems, for example in relation to invasibility, assembly and dynamical stability (Maynard et al., 2020).

Using our framework, we are able to obtain quantitative predictions for coexistence and abundances of an arbitrary set of species both in- and out-of-sample, even for data sets that comprise just a fraction of the possible sub-communities that may be formed from a fixed pool of species. Here, our simpler models are used to fit the relatively sparse data sets when fitting the full model is impossible. As the number of combinations that can be formed from a set of *n* species grows exponentially with *n*, this ability to predict species abundances out-of-sample is critical for exploring larger systems. Additionally, the good performance out-of-sample indicates that the models are capturing meaningful information about species interactions, rather than merely over-fitting the data.

Remarkably, we find that a model where interspecific interactions are approximated by a simplified structure, while intraspecific interactions are modeled more freely, achieves a comparable level of accuracy to the full model for the data sets. These results suggests that interspecific interactions in these systems are not completely idiosyncratic, but rather are largely characterized by a lower-dimensional structure. This observation agrees with the finding that simple rules govern the structure of interactions in microbial communities (Friedman et al., 2017), as well as recent work suggesting that plant-plant competitive interactions are characterized by low dimensionality (Stouffer et al., 2021). Similarly, these patterns are consistent with the success of a sparse-modeling approach demonstrating that, when considering only a few focal species of plants, many heterospecific interactions can be captured by a generic interaction term (Weiss-Lehman et al., 2022). Our results suggest that effective interspecific interactions sit in between extremely low-dimensional (i.e., as in the model with identical interspecific interactions, which has a poor performance across all data sets) and fully-structured pairwise interactions (arbitrary *B*).

Naturally, to develop these simple models we need to make certain strong assumptions about the community dynamics. Here we have ruled out “true-multistability,” in which a set of species can coexist at distinct configurations of abundances. We have also neglected higher-order or highly-nonlinear interactions, which would make the effect of species *i* on *j* context-dependent. While we would expect this framework to perform poorly when these assumptions are violated, the good agreement with experimental data suggests that departures from these strict assumptions are modest. However, relaxing these assumptions could further expand the applicability of these methods.

Another area that deserves further exploration is quantifying uncertainty in the point estimates produced by this approach. While one could gauge these effects via bootstrapping of experimental data, we instead advocate a fully Bayesian approach to uncertainty quantification, for example as implemented by Maynard et al. (2020), because deriving a posterior distribution for the matrix *B* would also allow one to determine the probability of coexistence for a set of species, and better characterize the correlations between species abundances. Both bootstrapping and MCMC would require evaluating the predicted values for a set of parameters a large number of times. In this respect, the computational gains afforded by our simplified models could be key to make such approaches viable in future studies.

The main outstanding problem with our approach is the assumption that dynamics have reached a steady state, and thus the observed community composition is the “true” final composition for the system. Violations of this assumption can greatly complicate inference. Suppose, for example, that we have two species, *A* and *B*, and that *A* excludes *B* asymptotically. If we sample this system before the extinction of *B*, we force the model to find parameters consistent with the robust coexistence of *A* and *B*, thereby biasing the results considerably. This problem of “spurious coexistence” is especially troublesome for microbial communities, where species’ presence is frequently determined by sequencing. Sequencing-based methods often detect some species at very low densities (Venturelli et al., 2018), potentially spuriously, making it difficult to discriminate between coexistence of rare species and actual extinctions masked by “background noise.” A Bayesian approach could make it possible to simultaneously impute the “true” final composition (i.e., which species are truly coexisting in the sample, taken as a latent parameter) as well as determine the distribution of parameters.

This general framework can be further extended in a number of directions. For example, one could introduce a more sophisticated error model assuming that species abundances are correlated within communities (e.g., overestimating the density of a predator could be associated with an underestimate of the densities of its prey). Similarly, we could assume more generally that observed densities 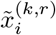 are sampled from a particular distribution (e.g., Gamma, Inverse Gaussian), with mean 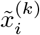 and ancillary parameters controlling the shape of the distribution. In this case, instead of minimizing the sum of squared deviations, we would maximize the likelihood of the parameters for the chosen distribution. Preliminary results shown in Supporting Information S9 demonstrate that indeed we are able to recover the parameters used to generate simulated data with specified error models. The ability to model the data using a variety of distributions would bring this framework one step closer to the flexibility that characterizes Generalized Linear Models, while maintaining the connection to ecological dynamics.

Overall, the methods presented make it easier to contrast experimental data with simple models for population dynamics, returning parameters that have clear ecological interpretation, and allowing us to test predictions in a straightforward manner. The type of ecological data examined here is appearing with increasing frequency in the ecological literature, and these methods provide a complementary (or alternative) approach to model-fitting via time-series analysis. The minimal data needed to fit the simplified models and the fact that each experimental community can be measured just once (possibly destructively) make this framework especially appealing and cost-effective.

## Acknowledgments

This work was supported by the National Science Foundation, award DEB-#2022742 to SA. We thank P. Lechon-Alonso and K. Della Libera for comments. We thank an anonymous reviewer and D. Stouffer for constructive criticism.

## Authorship statement

All authors contributed to the development of the methods. AS and SA wrote the code, analyzed the data, and wrote the manuscript. All authors edited the manuscript.

## Data accessibility statement

All data and model code necessary to recreate the analyses presented in this manuscript are archived at https://github.com/StefanoAllesina/skwara_et_al_2022

## Supporting Information

### S1. Description of the four data sets

#### Plant communities from Kuebbing et al. (2015)

The authors selected 8 phylogenetically-paired plant species (4 natives and 4 non-natives) typical of old-fields in the southeastern United States of America. The goal was to test how species richness affects seedling establishment and productivity below- and above-ground, and how the effects differ between native and non-native assemblages. The authors performed a nearly full-factorial design with richness varying from one to four species, performing 14 out of the 15 possible combinations. Each community had 20 replicates and within each replicate there were 12 individuals. We used the data from their biomass assay, in which they randomly selected 10 replicates from each treatment (native versus non-native) and estimate the biomass of all species. The remaining 10 replicates were used for the seedling establishment experiment (data not used here). The species used in each treatment are reported in Table S1.

**Table S1:** Species selection for the experiments by Kuebbing et al., 2015.

| Family | Native species | Non-native species |
| --- | --- | --- |
| Asteraceae (as) | *Achillea millefolium* | *Leucanthemum vulgare* |
| Fabaceae (fa) | *Lespedeza capitata* | *Lespedeza cuneata* |
| Lamiaceae (la) | *Pycnanthemum virginianum* | *Prunella vulgaris* |
| Poaceae (po) | *Sorghastrum nutans* | *Phleum pratense* |

#### Phytoplankton communities from Ghedini et al. (2022)

The authors cultured assemblages derived from a pool of five phytoplankton species (*Amphidinium carterae* [A], *Tetraselmis* sp [T]., *Dunaliella tertiolecta* [D], *Tisochrysis lutea* [Ti] and *Synechococcus* sp. [S]). They grew each species in isolation (3 replicates), two species together in all possible pairs (10 combinations, 3 replicates each), and all species together (5 replicates) for 10 days (corresponding to approximately ten generations). Time series for these data are reported in Figure S1. Samples were measured every other day. Here we analyze the data for day 8, because some of the species show a marked declined/increase on day 10.

#### Bacterial communities from Chu et al. (2021)

The authors performed experimental evolution by growing a strain of *Pseudomonas fluorescens* (P) along with all possible combinations of four strains, *Achromobacter* sp. (A), *Ochrobactrum* sp. (O), *Stenotrophomonas* sp. (S), and *Variovorax* sp. (V). The bacteria were grown in six replicates for each community in liquid media. Importantly, these strains form distinct morphological colonies, allowing the authors to estimate the density of each strain by plating an appropriately diluted sample and counting the number of Colony-Forming Units. The density of each species was determined at each transfer to fresh medium (transfers were performed every 7 days, Figure S2). Because *Variovax* sp. went extinct in the majority of communities and replicates, here we analyze the sub-communities measured for experiments that did not include *Variovax* sp.

**Figure S1:**
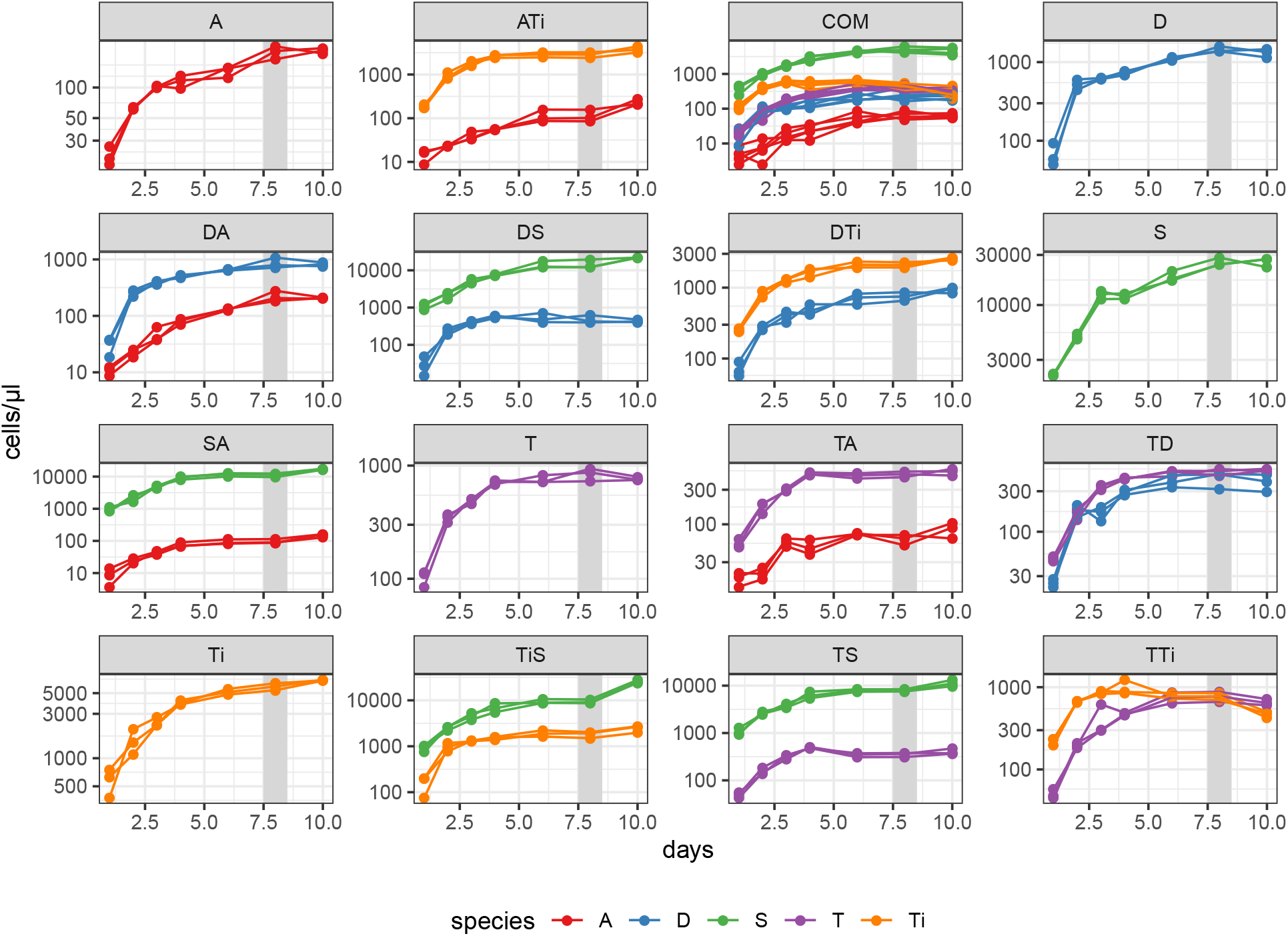
Time series for the growth of the 16 communities considered by Ghedini et al., 2022. For the analysis presented here, we considered the densities measured on day 8 (highlighted in the panels).

**Figure S2:**
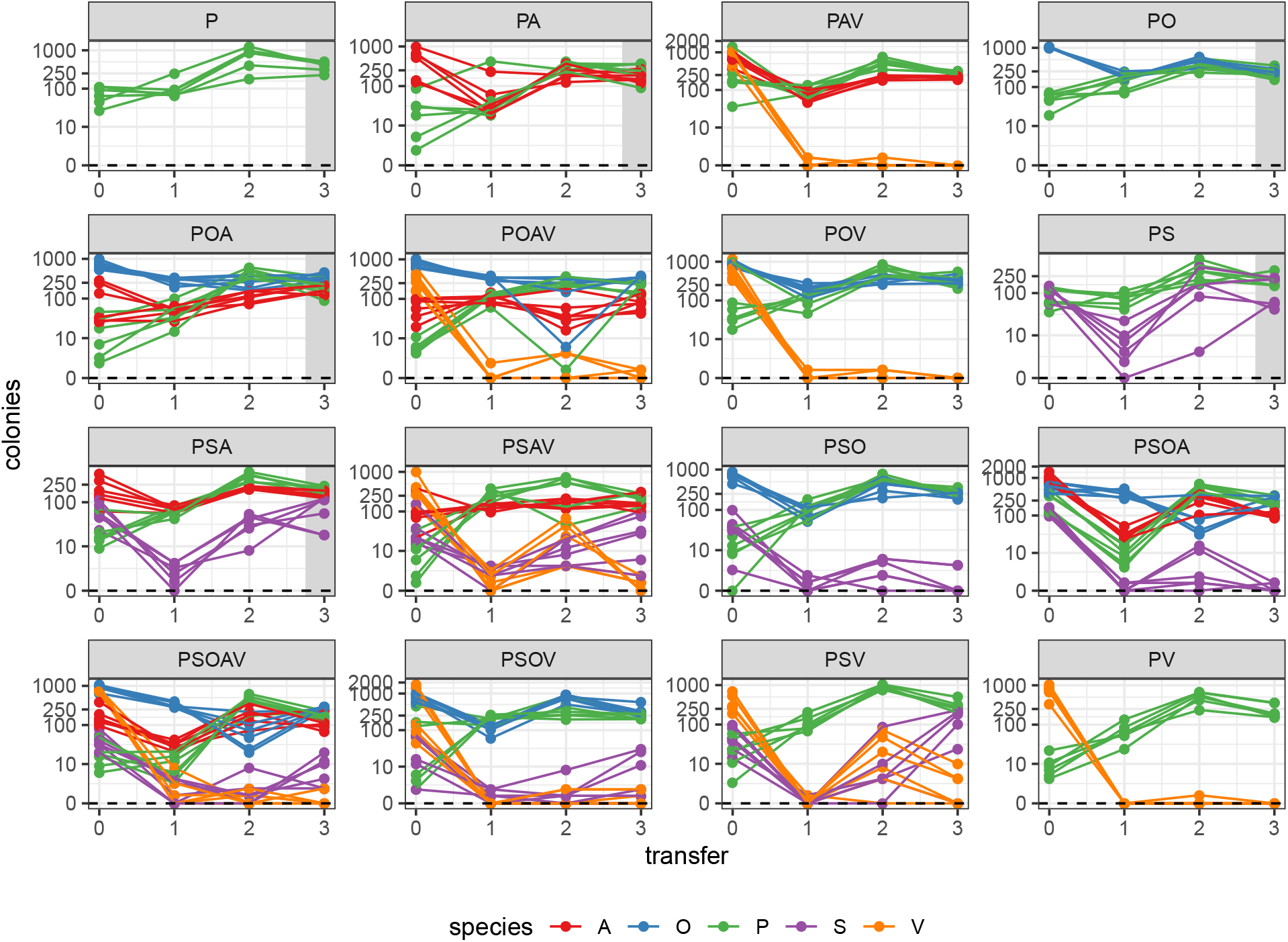
Time series for the growth of the 16 communities considered by Chu et al., 2021. For the analysis presented here, we considered the densities measured at the end of the experiment, for communities that did not include *Variovax* sp., because the strain went extinct in the majority of experiments. We also excluded communities where some of the replicates resulted in the extinctions of some of the species (PSO, PSOA). The points used for the analysis are highlighted in the panels.

### S2. Rank conditions for fitting all parameters

In order to fit the *n*^2^ parameters of the interaction matrix *B* in our model, we need to have observed a sufficiently large variety of communities. For simplicity, suppose that we are simulating data from the model as specified by a given matrix *B*, which details the interactions between the *n* species in the pool. For each possible set of species *k* (i.e., all single species, all possible pairs, etc.), we compute the solution 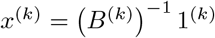 and, if all components of *x*^(*k*)^ are positive, we record them in a row of matrix *X*, along with zeros with all the species that are in the pool but not present in community *k*. We ask when we can recover all the coefficients in *B* from *X*.

Under these conditions, to recover the coefficients of the matrix *B* we need to be able to find a unique solution for *BX^T^* = *P^T^*. We can write this as:

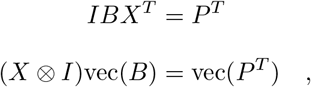

where vec(*A*) is the vectorization operator (stacking all the columns in a matrix into a vector), and ⊗ is the Kronecker product. Note that (as explained in details in Supporting Information, S4), we cannot determine all elements of *P* without having access to matrix *B*. We can however be certain that whenever *X_ij_ >* 0, then *P_ij_* = 1. If we then call 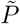 a matrix that has 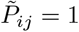 whenever *X_ij_ >* 0 and 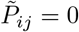 elsewhere. We can rewrite the system of equations above as:

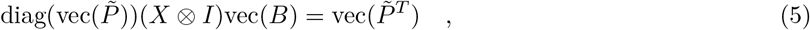

where diag(*b*) creates a diagonal matrix with vector *b* on the diagonal. It is now apparent that solving the matrix equation for the *n*^2^ entries of *B* requires the matrix 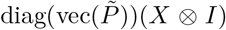 to have rank *n*^2^. This condition will be met only when each of the *n* species in a system are present in at least *n* experiments, and each pair of species appears in at least one experiment. Note that if any two species were to always co-occur alongside each other in the experimental data, we would not be able to fit the full *B* matrix—in this case, we would be unable to distinguish the effects of the two species from one another.

The procedure is the same for observed data; all we need to do is to substitute matrix *X* in Eq. 5 with the matrix of observed densities obtained by retaining a single row for each community (i.e., removing replicate experiments). These conditions are equivalent to those found in Maynard et al. (2020).

### S3. Naïve approach

Here we rewrite the naïve approach of Maynard et al. (2020) in matrix form, thereby showing that it does not minimize the SSQ and therefore does not yield the m.l.e. for matrix *B*.

Taking the equation Eq. 4, and using the matrix 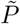 as above, we find:

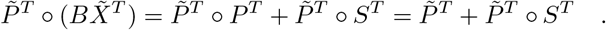

where ∘ is the element-by-element (Hadamard) product.

If we proceed as above and attempt to solve the system of equations

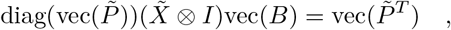

for the matrix *B* by taking the Moore-Penrose pseudoinverse,

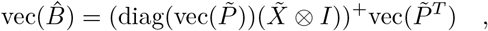

where *A*^+^ = (*A^T^ A*)^−1^*A^T^*, what we would be minimizing is 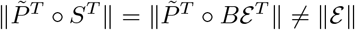. As such, this method will not be the m.l.e. for *B* for any case in which the measurements are noisy, even if the process generating the data would follow the model precisely.

### S4. Connections to the Generalized Lotka-Volterra model

The Generalized Lotka-Volterra (GLV) model is arguably the simplest nonlinear model for population dynamics, and has been studied for more than a century (Lotka, 1920; Volterra, 1926). It can be written as:

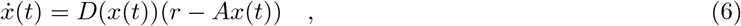

where *ẋ*(*t*) = *dx*(*t*)/*dt*, *x*(*t*) is a column vector detailing the densities of all species at time *t*, *r* is a vector of intrinsic growth/death rates, *A* is a matrix of species’ interactions, and *D*(*x*(*t*)) is a diagonal matrix with *x*(*t*) on the diagonal. We can divide each element of *A* by the corresponding growth rate, obtaining:

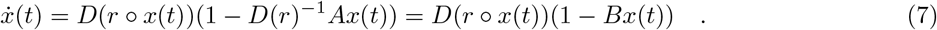

These equations hold whenever all species are present, or when we initialize the system using a subset of species *k*. The model admits up to 2^*n*^ equilibria, of which only one can have all positive components (feasible coexistence equilibrium), and the others are “boundary” equilibria, in which a certain subset of species *k* can coexist, and the remaining species are absent. For each subset of species *k*, a feasible equilibrium— when it exists—is found solving 1^(*k*)^− *B*^(*k*)^*x*^(*k*)^ = 0^(*k*)^, i.e., 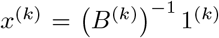, which is exactly our Eq. 3. Therefore, as already noted by Maynard et al. (2020), fitting the statistical models presented here is equivalent to finding a GLV model with an equilibrium structure that is as close as possible to the observed data. Note that this also makes clear that, using “static” observations of several sub-communities, we cannot find *A* and *r* separately, only the composite parameters *B* = *D*(*r*)^−1^*A*.

In this context, the predicted matrix *X* (for a given matrix *B*) is simply a collection of feasible equilibria for some of the sub-systems we can form from the pool of *n* species. Take one of the rows of *X*, 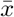, containing the density of the |*k*| species forming the feasible equilibrium, along with zeros for the species that are absent. Because of the equilibrium condition, for all 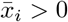, we have

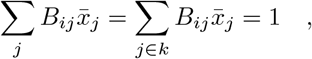

for the remaining species (i.e., the species *x_i_* not present in *k*), we have

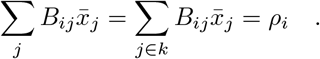

The quantity *ρ_i_* can be interpreted as the effect of the “resident” species (i.e., those for which 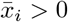) on the growth rate of the invader *i* when entering the community at low abundance. Write the per-capita growth rates:

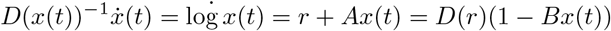

If the system is resting at 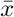, we have that whenever 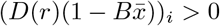, species *i* has positive growth rate when invading the system at low abundance. If we assume that the growth rates are positive for all species, we have that the inequality for species *i* is satisfied whenever 1 − *ρ_i_ >* 0, and therefore *ρ_i_* < 1. If the sign of *r_i_* is negative, the inequality is reversed.

Therefore, when we compute *P* = *XB^T^*, we have that *P_ij_* = 1 whenever the species *j* is part of the community specified by row *i* of *X*, and that (assuming *r_i_ >* 0) whenever *P_ij_* < 1 for a species that is not part of the community, then the species can invade the community when rare, and cannot invade whenever *P_ij_ >* 1. If moreover we have that the species in *k* are resting at a stable equilibrium, then any row of *P* for which *P_ij_* ≥ 1 for all species represents a feasible, stable and non-invasible equilibrium for the system, also called a “saturated rest point” (Hofbauer et al., 1998; Serván et al., 2018). Serván & Allesina (2021) showed that these equilibria are also final configurations for the assembly paths one can form from the pool of *n* species.

### S5. Derivation of the simplified models from a consumer-resource framework

Having established a parallel between our statistical model and the equilibrium structure of a Generalized Lotka-Volterra model, we use this equivalence as a jumping board to derive simplified versions of our model. In particular, we consider a simple MacArthur’s consumer-resource model (MacArthur, 1970), in which we have *n* consumers and *m* resources (assuming *m* ≥ *n*, a necessary condition for the coexistence of the *n* consumers):

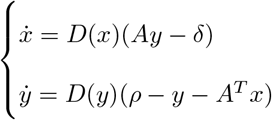

where *ẋ* = *dx*(*t*)/*dt* and *ẏ* = *dy*(*t*)/*dt* are vectors describing the change in the density of the *n* consumers (stored in *x*) and the *m* resources, (*y*), respectively. The vector *δ* stores the death (growth) rate of the consumers, *ρ* that of the resources, and the *n* × *m* matrix *A* the attack rates of consumers on resources; *D*(*x*) defines a diagonal matrix with *x* on the diagonal. If we assume that the dynamics of the resources are fast compared to those of the consumers, we can solve for the density of the resources attained for a given state of the consumers:

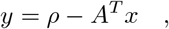

and substitute this value in the equations for the consumers, obtaining:

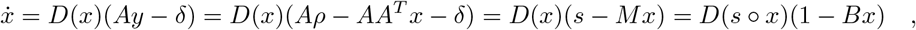

where we have defined *s* = *Aρ* − *δ*, *M* = *AA^T^* and *B* = *D*(*s*)^−1^*M*. This is a Generalized Lotka-Volterra model, and the term in parenthesis is exactly what we have considered for our statistical model.

In principle, the matrix *A* can detail the interactions with an arbitrary number of resources *m* ≥ *n*. To simplify the statistical model and reduce the number of free parameters, we assume a particular structure for *A*. Suppose that each consumer has access to private resources, as well as a shared resources. In the simplest case, with one private resource for each consumer and one resource shared by all consumers, the matrix *A* has the following structure:

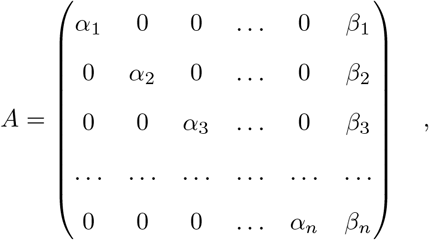

where the vector *α* collects the attack rates for the consumers on their private resources and the vector *β* the attack rates for each consumer and the shared resource. Then we have that *M* = *AA^T^* is simply:

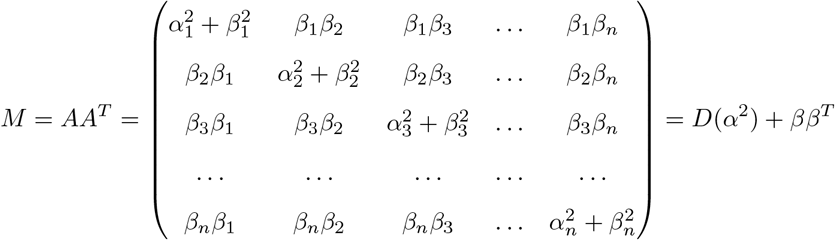

Dividing each row of *M* by the corresponding *s_i_*, we obtain:

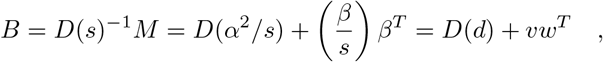

where *d* = *α*^2^/*s*, *v* = *β/s* and *w* = *β*. As such, in this case we can fully define the matrix *B* using 3*n* − 1 parameters (with the −1 due to the fact that elements *v_i_w_j_* always appear as products). We can complicate this model by adding extra shared resources, each time adding further parameters. In particular, if we wanted to model off-diagonal (i.e., inter-specific interactions) as the sum of two rank-1 matrices, we could write:

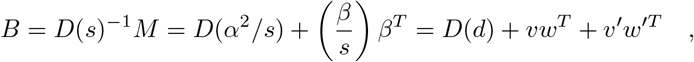

adding only 2*n*−3 parameters; in fact, without loss of generality we can assume that *v^T^ v*^′^ = 0 and *w^T^ w*^′^ = 0, i.e., the two left vectors are orthogonal and the two right-vectors are also orthogonal. The orthogonality condition allows us to solve for one of the parameters in each vector. Adding another rank-1 matrix, say *v*^′′^*w*^′′*T*^, adds a further 2*n* − 5 parameters, and so on.

Can the model be simplified further? If we assume that all the *s_i_* are the same, then the matrix can be rewritten as *B* = *D*(*d*) + *vv^T^*, and is therefore symmetric (2*n* parameters); if further we assume that all consumers have the same attack rate on the resources, the matrix can be written as *B* = *D*(*d*) + *α*11^*T*^ (*n* + 1 parameters).

We have shown how the consumer-resource framework can be used to derive simplified versions of the statistical model, thereby lifting some of the requirements on the data (see section S7 below). While in a consumer-resource model we would expect all attack rates to be positive, we can relax this condition to have a more general and flexible model.

### S6. Fitting the simplified models

The main computational hurdle one faces when fitting the original model with *n*^2^ parameters is the need to invert a sub-matrix of *B*, *B*^(*k*)^ for every species combination *k*, in order to compute the predicted values. Inverting matrices is computationally costly, requiring on the order of approximately *n*^2.4^ operations for an *n* × *n* matrix, even using the best available algorithms.

This problem is greatly reduced for the simplified models, as we can easily obtain a closed-form analytical expression for the inverse of a sub-matrix by applying the Sherman-Morrison formula(Sherman & Morrison, 1950). In particular, if we have a matrix:

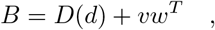

for arbitrary vectors *d*, *v* and *w*, we can rewrite it without loss of generality as:

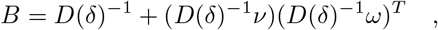

by defining *δ* = 1/*d*, *ν* = *v/d* and *ω* = *w/d* and assuming element-by-element (Hadamard) division. With this notation, the inverse of *B* becomes:

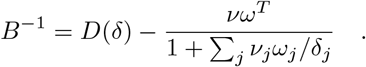

The predicted abundance for the species in the community can be computed as:

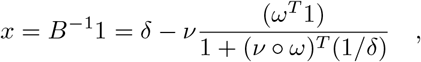

Even more conveniently, the predicted abundance for a sub-community be readily computed. Simply define a vector of presence/absence *π_k_* such that *π_ki_* = 1 if species *i* is part of the sub-community and 0 otherwise. Then, we can write *x*^(*k*)^, containing the predicted density of the species when present, and 0 otherwise, as:

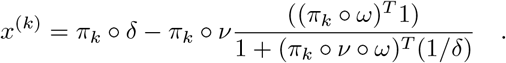

Similar (and even simpler) calculations can be performed for the further simplifications of the model. Having access to a linear-time, analytical way to compute the predicted densities speeds up the calculation enormously, allowing our methods to be applied to larger sets of experiments.

### S7. Data requirements to fit the simplified model

As detailed in Appendix S2, to fit the full model we need to identify the *n*^2^ coefficients of the matrix *B*. This can be accomplished by writing a set of equations based on the observed data. In particular, having observed a certain community *k* in which *m* species coexist, we can write *m* equations. For example, having observed species one growing in monoculture (labeled community 1), we can write:

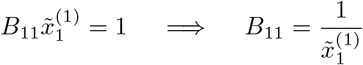

Similarly, when we have observed species one and two growing together (labeled community 2), we can write two equations:

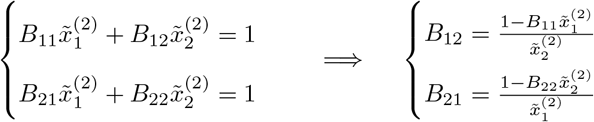

As such, having observed all monocultures we can write *n* equations, and having observed all the 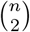 pairs we can write *n*(*n* − 1) equations—for a total of *n*^2^ equations, allowing us to solve for all the coefficients of *B*. Naturally, the same can be accomplished when we have observed all the species growing together (yielding *n* equations), as well as all the leave-one-out communities (*n* communities, yielding *n* − 1 equations each), or any other combination of experimental observations that allows us to write *n*^2^ linearly independent equations.

Take the simplest of the reduced models presented in section S5 and S6:

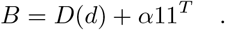

Following the same logic, it is obvious that having observed all the monocultures (yielding 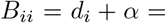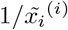) and any of the pairs (allowing us to solve for *α*) would be sufficient to parameterize the model.

The more interesting and complicated case is that of the most complex of the reduced models: *B* = *D*(*d*) + *vw^T^*. Here we show that at a minimum we need to be able to write 3*n* − 1 independent equations, in order to solve for the *n* diagonal coefficients of *B* and 2*n* − 1 off-diagonal coefficients of *B*. Moreover, we demonstrate that not all combinations of off-diagonal coefficients identify the parameters uniquely, and provide a simple test to determine the identifiability of the parameters.

Because the method to identify the parameters is slightly more complex and abstract than in the case of the full model, we illustrate it with an example. Take the four-species community defined by the parameters:

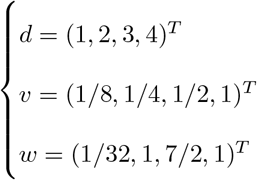

Yielding the matrix *B*:

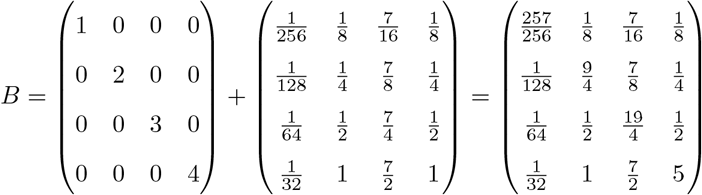

This system allows for the coexistence for all combinations of species. For simplicity, in the examples below, we will only use the “observed” data for monocultures and pairs:

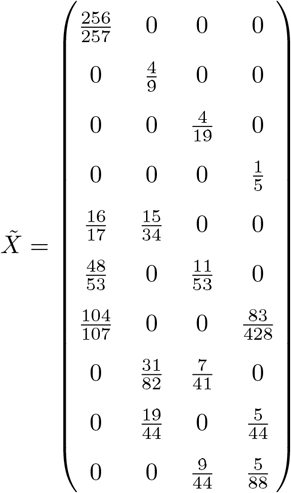

Clearly, if we had access to all the rows of 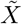, we would be able to infer an arbitrary *B* (as in the full model), and therefore also any *B* with the special form *B* = *D*(*d*) + *vw^T^*. Our goal is to show that we do not need to have observed all the monocultures and pairs to infer the parameters of this simplified form.

Before we start, we note that *G* = *vw^T^* is defined uniquely by 2*n* − 1 (rather than 2*n*) parameters. In fact, we can take any *θ* 0 and define *v*^′^ = *θv* and 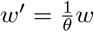 such that *G* = *vw^T^* = *v*^′^*w*^′*T*^.

Now consider the case in which we have observed all the monocultures, all the pairs involving species one, and the pair involving species two and three. I.e., we have observed all the monocultures, but only four pairs of the six potential. We can use this data to solve for the coefficients *B*_11_, *B*_12_, *B*_13_, and *B*_14_:

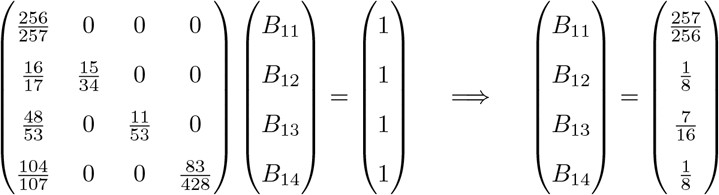

Using the appropriate observations, we can also identify *B*_22_, *B*_21_, *B*_23_, *B*_31_, *B*_33_, *B*_41_ and *B*_44_. In summary, we have identified 3*n* − 1 coefficients of *B*: the *n* elements on the diagonal, as well as 2*n* − 1 elements in the off-diagonal part of the matrix.

Note that *B_ii_* = *d_i_* + *G_ii_*, while *B_ij_* = *G_ij_* when *i j*. We have therefore observed a partial matrix *G*:

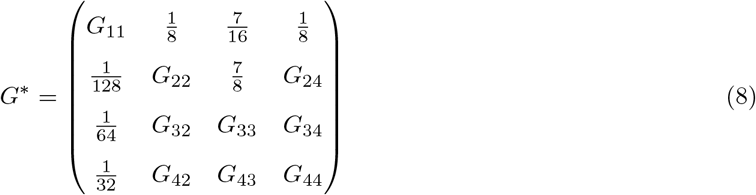

Is there a unique way to “complete” the matrix *G*, knowing that it must be rank-1? This problem is known in mathematics as the “matrix completion” problem, and has been studied extensively. In particular, a partially-specified matrix admits at least one rank-1 completion whenever it meets two conditions (Hadwin et al., 2006). The first condition, called the “zero row or column property” is very simple: if one of the specified coefficients is zero, then all the specified coefficients in either the corresponding row or column must also be zero. In our case, we have no zero coefficients, and thus the property is trivially satisfied.

The second condition is based on the bipartite graph 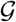 that we can construct by taking the labels for the rows and columns of *G*^*^, *r_i_* and *c_i_* respectively, as nodes and drawing an undirected edge between *r_i_* and *c_j_* whenever 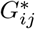 is specified. The second condition, called the “cycle property,” has to do with the consistency of the undirected cycles in the bipartite graph 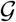. For the partially-specified matrix in Eq. 8, the graph has no undirected cycles (Fig. S3), and therefore the property is trivially satisfied. Finally, if the graph 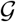 is connected (i.e., there is a path connecting any node to any other), the rank-1 completion is unique. Thus, by inspecting the graph corresponding to matrix *G*^*^ in Eq. 8, we conclude that a matrix completion of *G*^*^ exists and is unique. Having established that a completion exists and is unique, we then ask how we can find the coefficients needed to complete the matrix.

**Figure S3:**
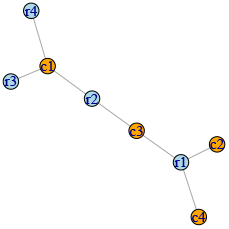
Bipartite graph associated with the partially-specified matrix in Eq. 8. The graph represents a connected tree, and as such a completion of matrix *G*^*^ exists and is unique.

For this final step, one can devise several alternative algorithms. One of the simplest approaches, from the conceptual point of view, is based on the fact that any *k* × *k* submatrix of a rank-1 matrix, obtained selecting a certain set of rows and corresponding columns, must also be rank-1. As such, the determinant of any submatrix of *G*^*^ with at least two rows and columns must be zero. We can therefore form equations to solve for the remaining coefficients. For example, considering the submatrices induced by the rows and columns {1, 2}, {1, 3}, {2, 3}, and {1, 2, 3}, we can write:

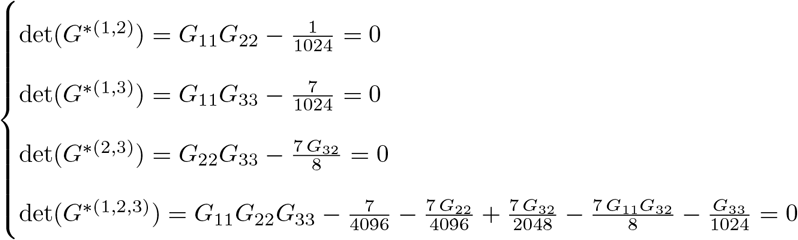

Yielding 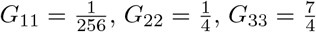 and 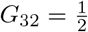. Having identified these coefficients, we write equations for det(*G*^*(1,4)^) to find *G*_44_ = 1, and then the equations for det(*G*^*(2,4)^) and det(*G*^*(1,2,4)^) to solve for 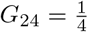 and *G*_42_ = 1. Finally, to complete the matrix *G*^*^, we solve the equations for det(*G*^*(3,4)^) and det(*G*^*(1,3,4)^), yielding 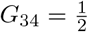 and 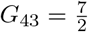.

In summary, provided with 2*n* − 1 nonzero coefficients of *G* satisfying the connectedness of the corresponding graph 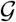, we can complete the matrix and the completion will be unique. Note that we are specifying only 2*n* − 1 coefficients, which implies that 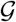 has only 2*n* − 1 edges. As such, if 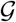 is connected, it is necessarily a tree. A tree automatically satisfies the cycle condition in (Hadwin et al., 2006), and if the coefficients are nonzero, connectedness is necessary and sufficient for the existence of a unique solution. Because the solution is unique, numerical methods could easily substitute the algebra we performed above.

Finally, having identified all of the coefficients *G_ii_*, and provided with the estimates of *B_ii_* we obtain *d_i_* by difference.

To demonstrate that not all choices of 2*n*−1 off-diagonal coefficients would satisfy the conditions for existence and uniqueness, consider the same system and use the monocultures and the experiments in which we grow species (1, 2), (2,3), (3,4) and (4,1) together. In total, we should be able to identify 3*n* coefficients of *B*, including all the diagonal coefficients and 2*n* off-diagonal coefficients. Excluding one of them arbitrarily (this has no effect on the result), we end up with the partially-specified matrix:

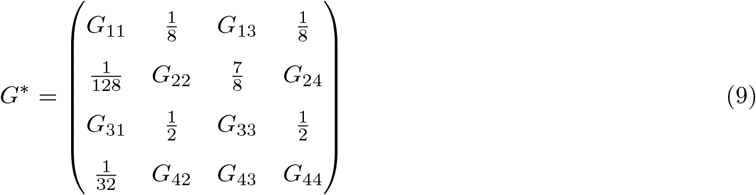

In Figure S4 we show that the induced graph 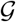 is not connected (and in fact, not a tree). As such, if the matrix were to satisfy the cycle condition (it does), we would have multiple solutions (in this case, infinitely many), while if it did not, we would have no possible completion.

**Figure S4:**
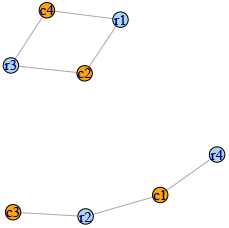
Bipartite graph associated with the partially-specified matrix in Eq. 9. The graph does not represent a connected tree, and as such a completion of matrix either does not exist, or is not unique.

In summary, to identify the parameters of the simplified model we need to have sufficient data to determine both the *n* diagonal coefficients of *B* as well as 2*n* − 1 off-diagonal coefficients, ensuring that the induced graph 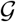 associated with the matrix *G*^*^ is connected. This condition is both necessary and sufficient to guarantee the existence and the uniqueness of a solution. Finally, because we need to identify *n*^2^ parameters (quadratic in *n*) for the full model, and only 3*n* − 1 (linear in *n*) for the simplified model,the benefits of the simplified model will be markedly greater for larger *n*.

### S8. Full results

#### Iterative Algorithm

The iterative method proposed in the main text improves over the naïve approach reported in section S2 in all cases (Figures S3, S4, and S5).

**Figure S5:**
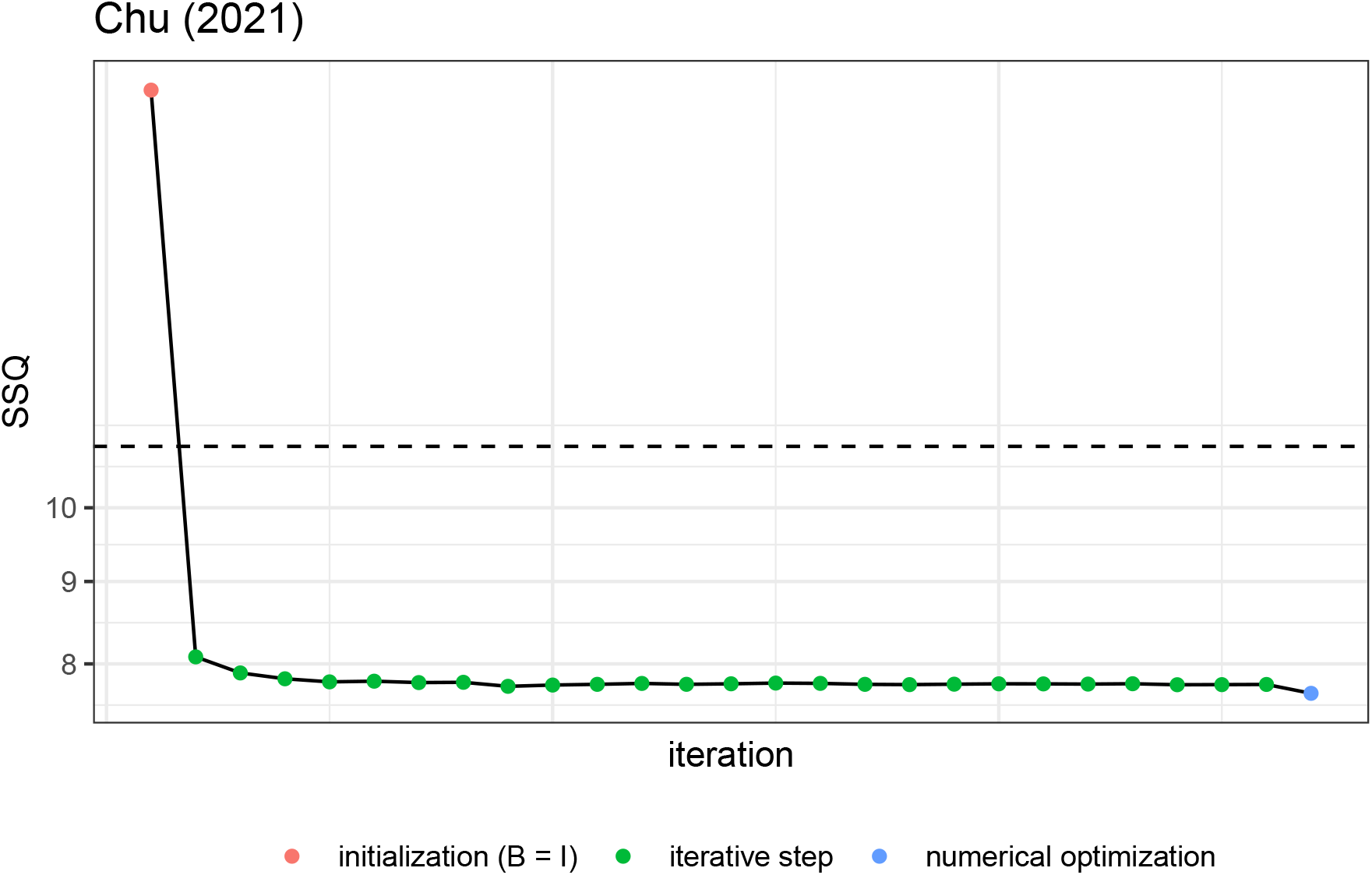
As Figure 1, but for the data of Chu et al., 2021.

**Figure S6:**
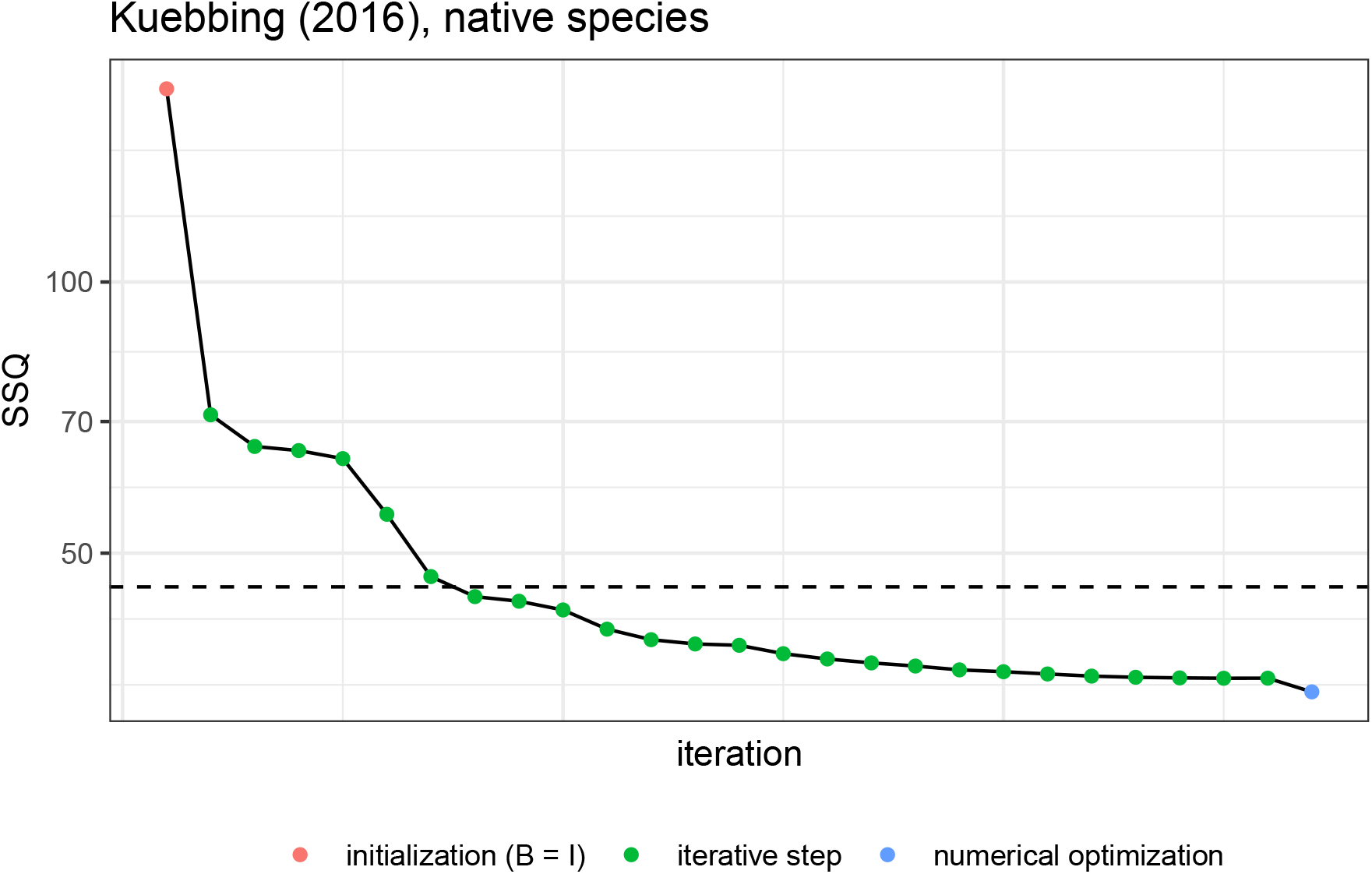
As Figure 1, but for the data of Kuebbing et al. (2015), native plants).

**Figure S7:**
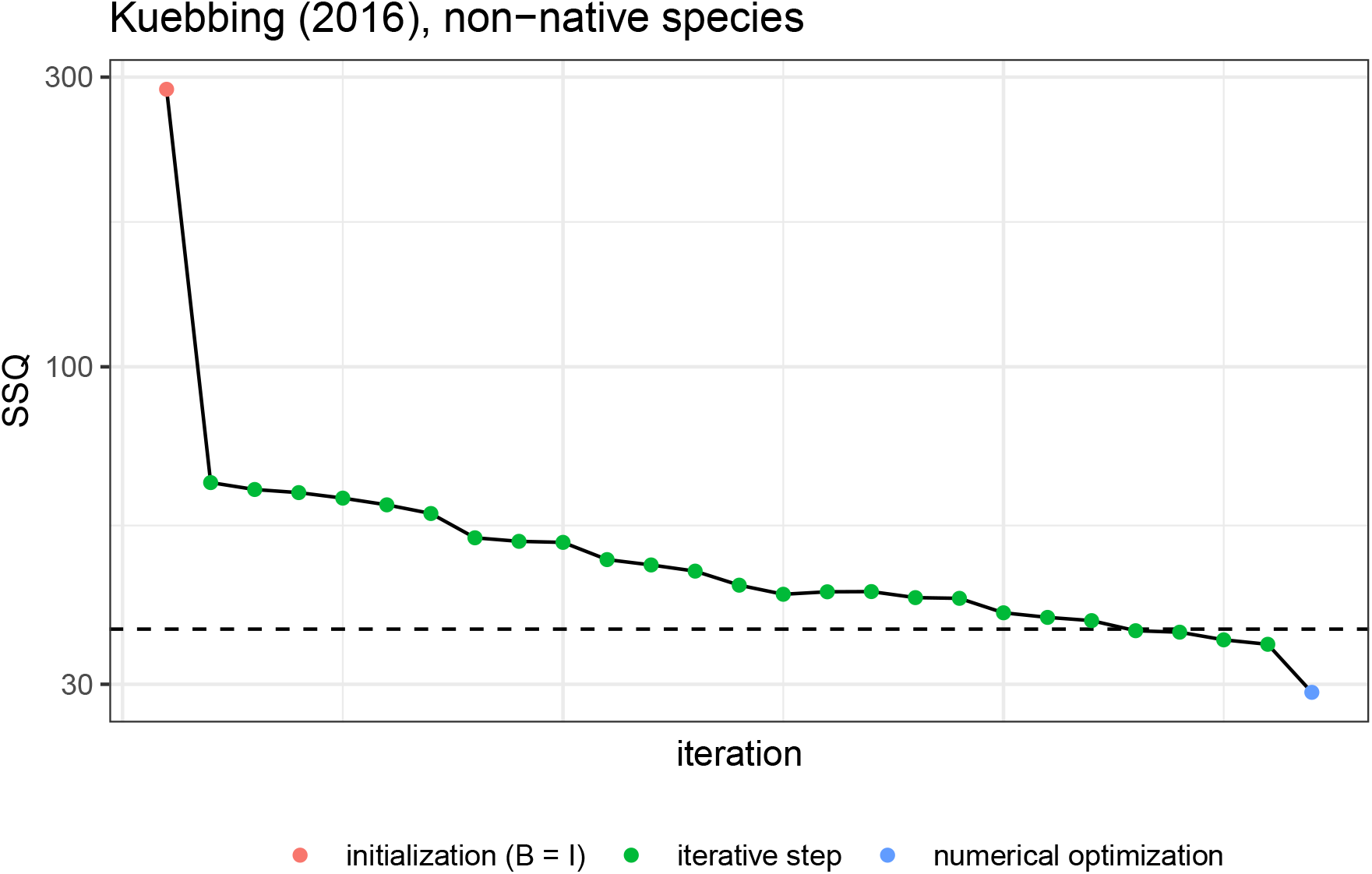
As Figure 1, but for the data of Kuebbing et al. (2015), non-native plants.

#### Model comparison

For all data sets, we find that the simplified model with 3*n* − 1 parameters, in which the interaction matrix is constrained to have form *B* = *D*(*d*) + *vw^T^* perform similarly to the full model with *n*^2^ parameters (Figures S6, S7, and S8).

**Figure S8:**
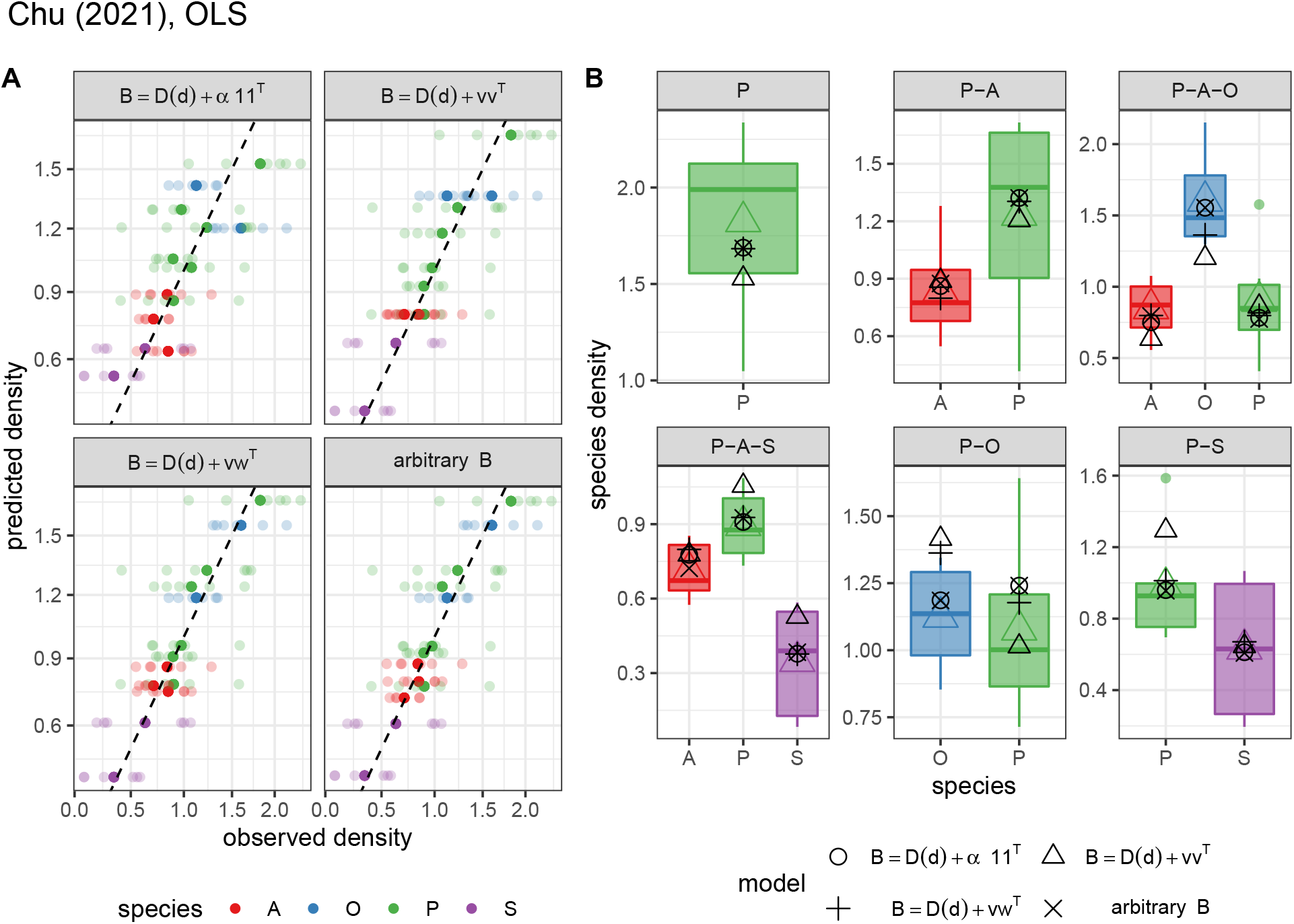
As Figure 2, but for the data of Chu et al., 2021.

**Figure S9:**
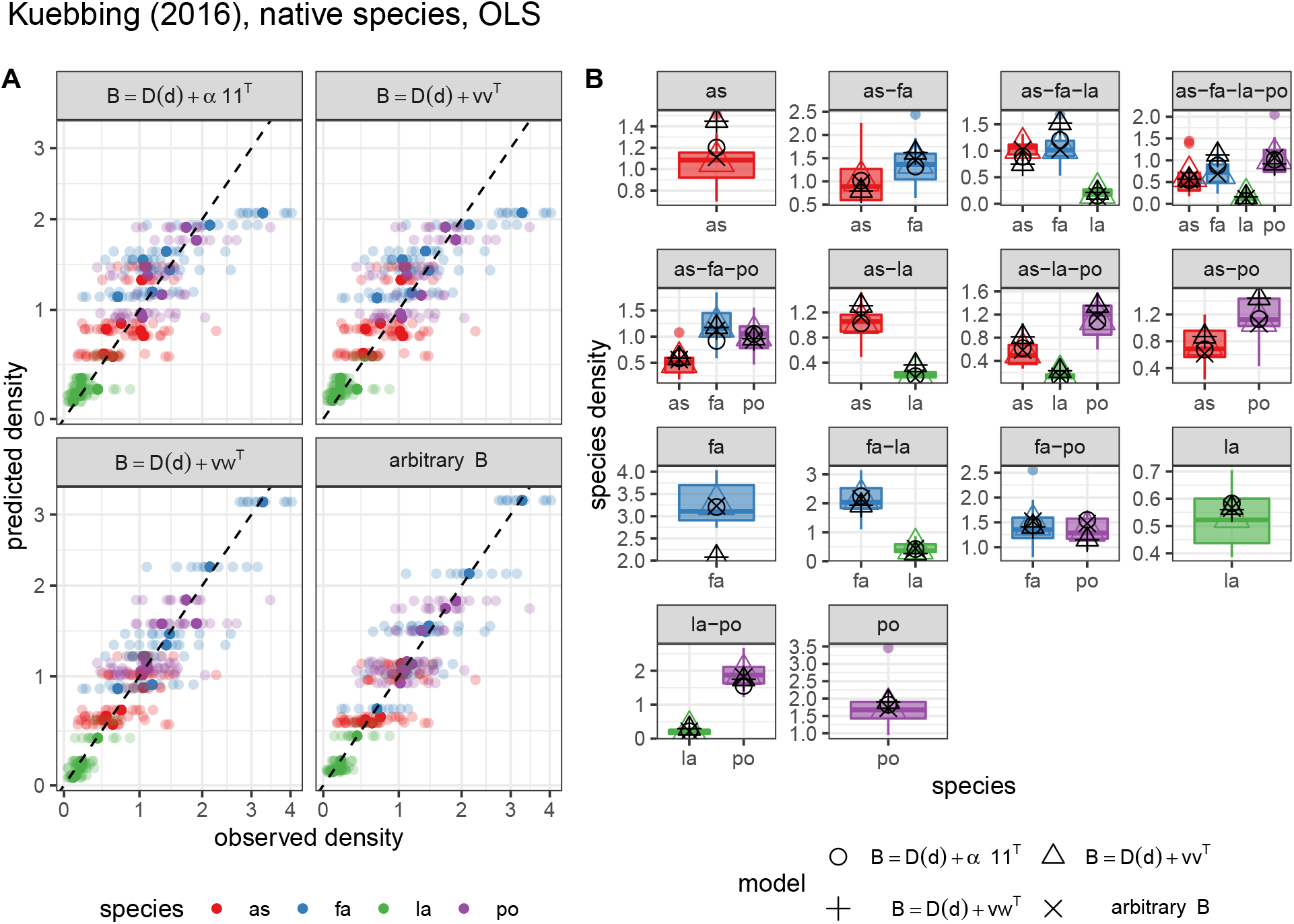
As Figure 2, but for the data of Kuebbing et al. (2015), native plants.

**Figure S10:**
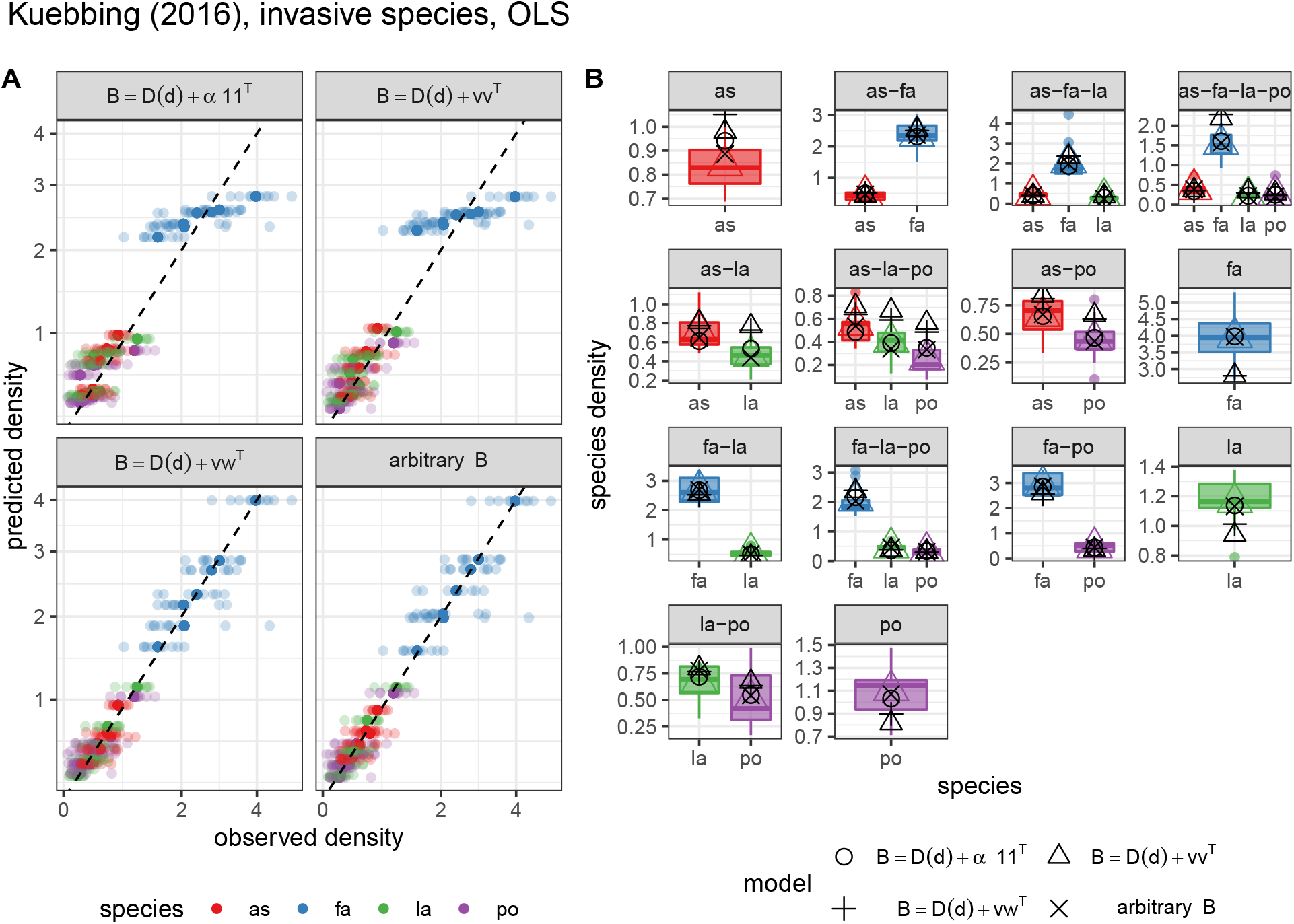
As Figure 2, but for the data of Kuebbing et al. (2015), non-native plants.

#### Ordinary Least Squares vs. Weighted Least Squares

When attempting to minimize the weighted sum of squared deviations, rather than the sum of squared deviations, we obtain a better fit for the species with low abundance, at the cost of a slightly less precise fit for the highly abundant species (Figures S9, S10, and S11).

**Figure S11:**
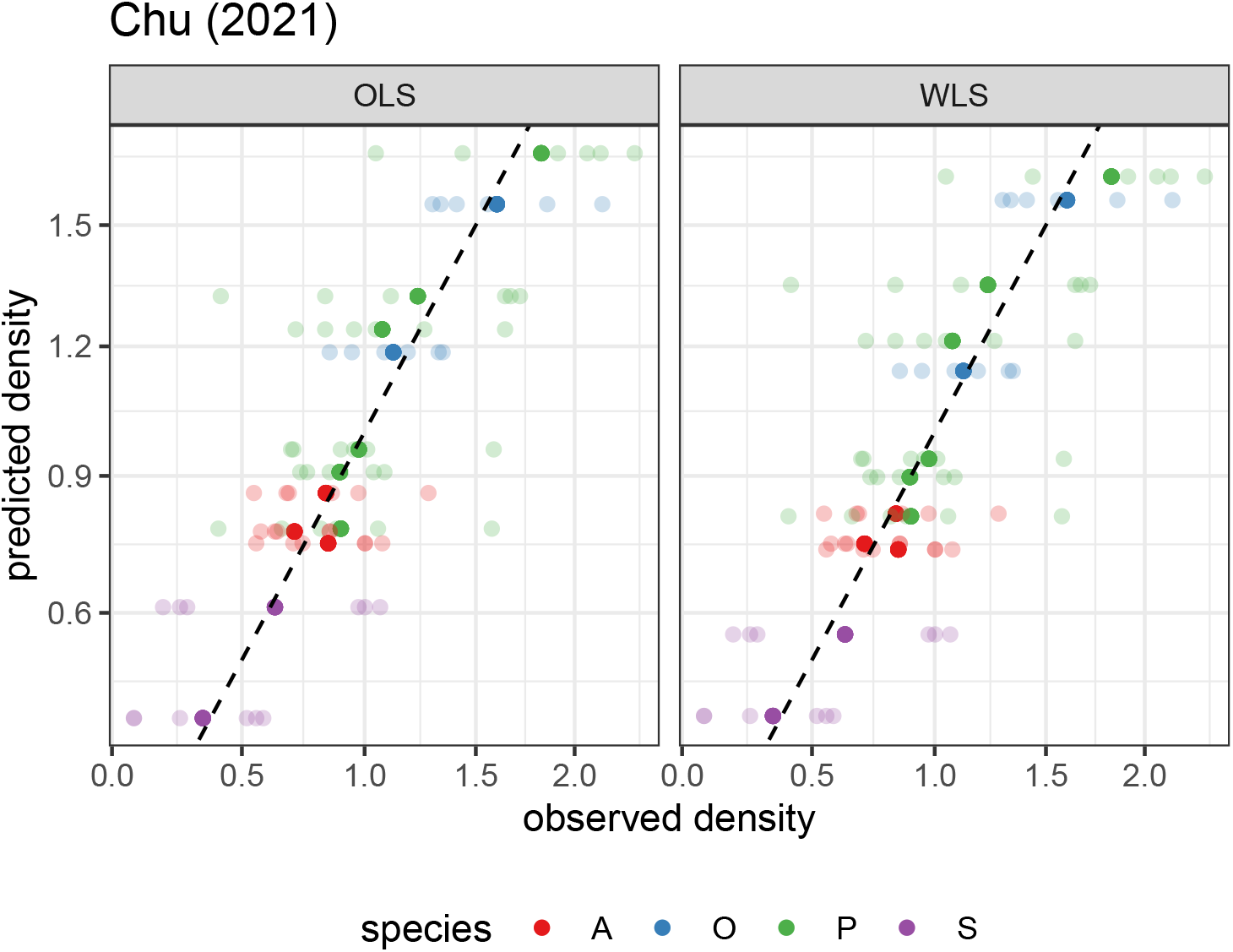
As Figure 5, but for the data of Chu et al., 2021.

**Figure S12:**
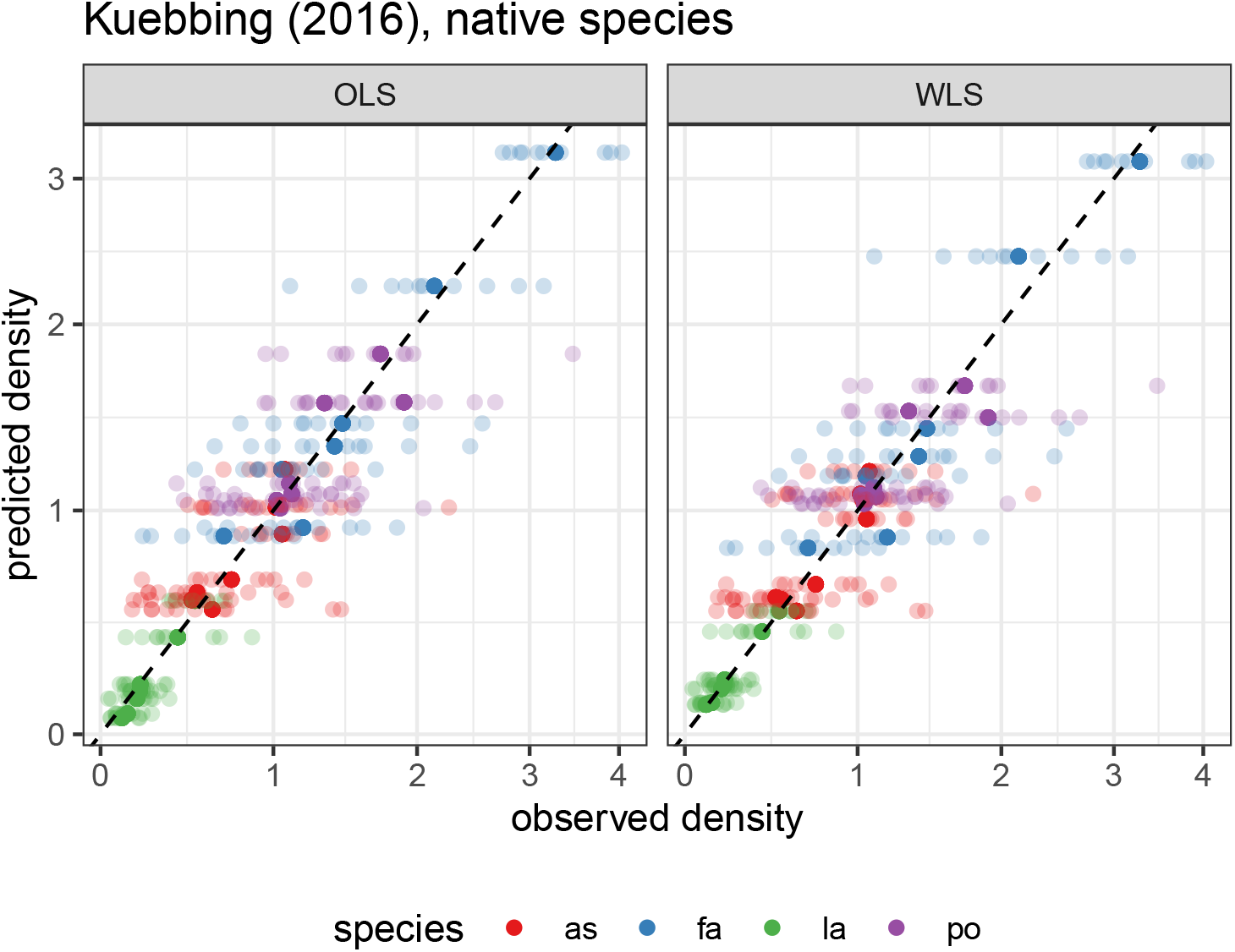
As Figure 5, but for the data of Kuebbing et al. (2015), native plants.

**Figure S13:**
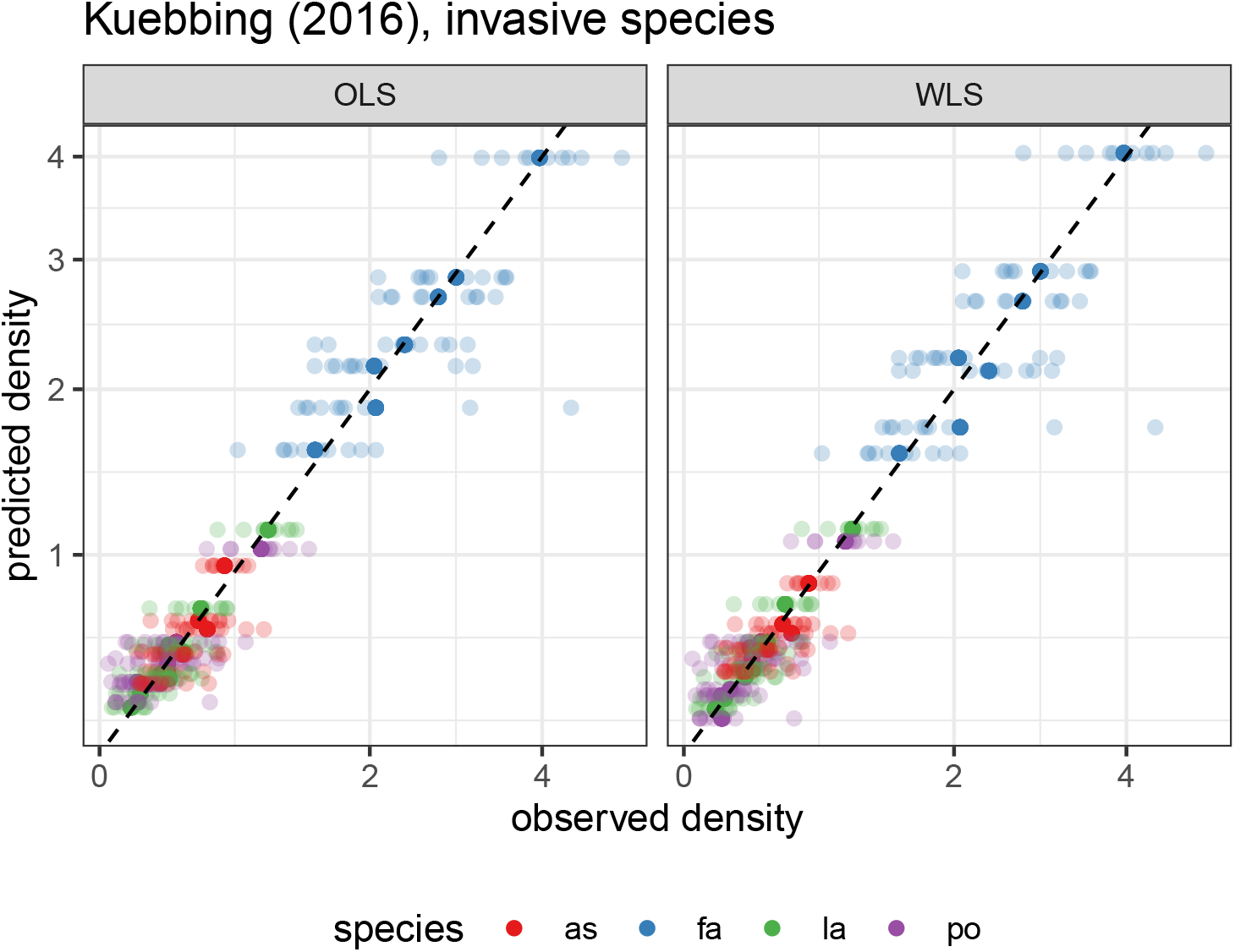
As Figure 5, but for the data of Kuebbing et al. (2015), non-native plants.

#### Out-of-sample predictions for the simplfied model *B* = *D*(*d*) + *vw^T^*

For the data sets of Kuebbing et al. (2015), even using the simplified model *B* = *D*(*d*) + *vw^T^* we obtain excellent out-of-sample predictions (Figures S13 and S14): we correctly predict that the species coexist in the observed communities, and predicted densities that match the observed values quite closely. For the data from Chu et al. (2021), on the other hand, we make one qualitatively incorrect prediction (lack of coexistence, when the community is observed to coexist empirically), and also the quantitative predictions are worse, especially for the *Ochrobactrum* sp., which appears in only two communities (Figure S12). In all cases, using OLS or WLS yields similar results.

**Figure S14:**
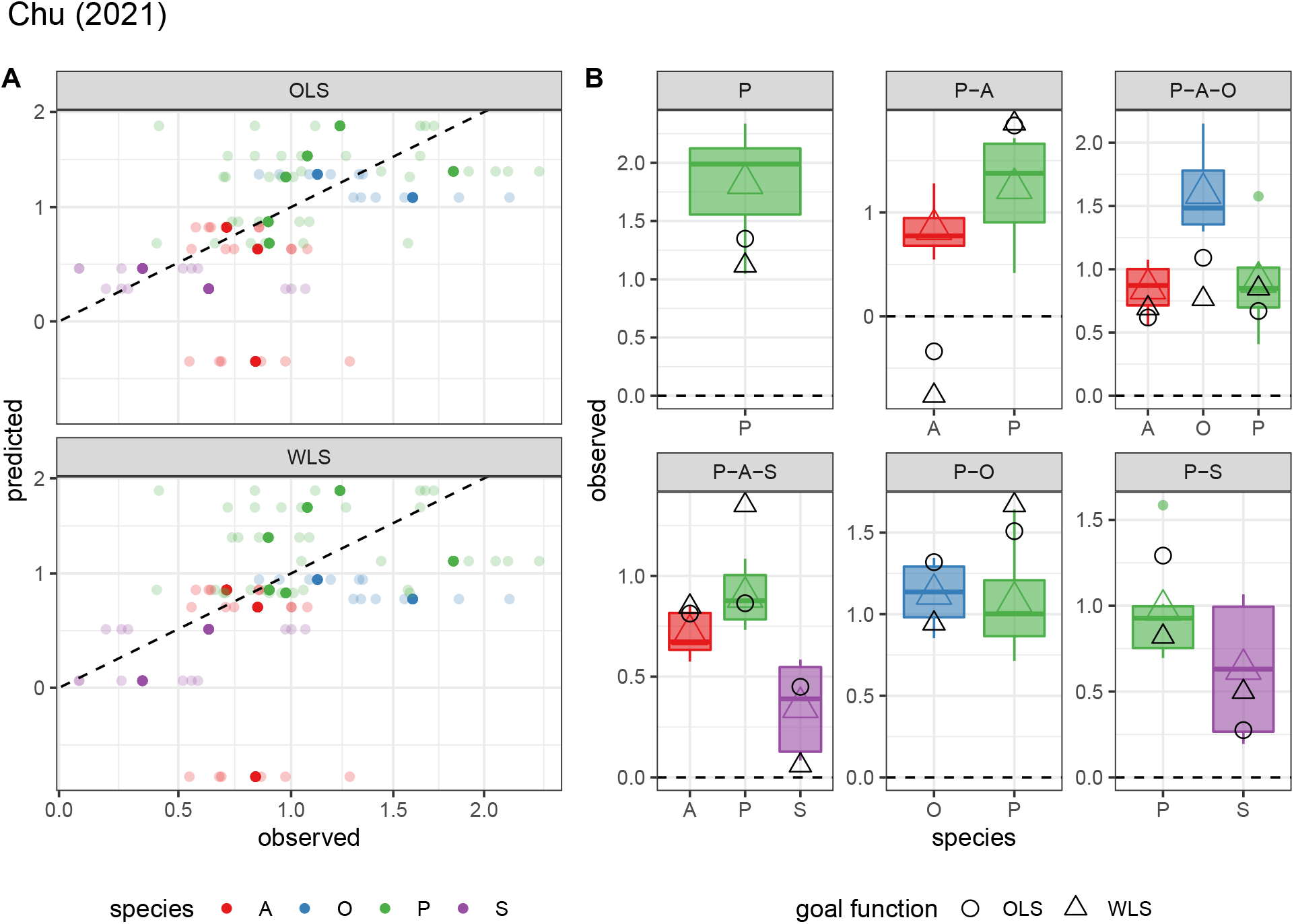
As Figure 6, but for the data of Chu et al., 2021. When we fit the model without the experiments in which *Pseudomonas fluorescens* and *Achromobacter* sp. are grown together, we obtain a qualitatively wrong prediction (lack of coexistence). *Ochrobactrum* sp. appears in only two communities, and when one of the communities is removed the fit is greatly reduced.

**Figure S15:**
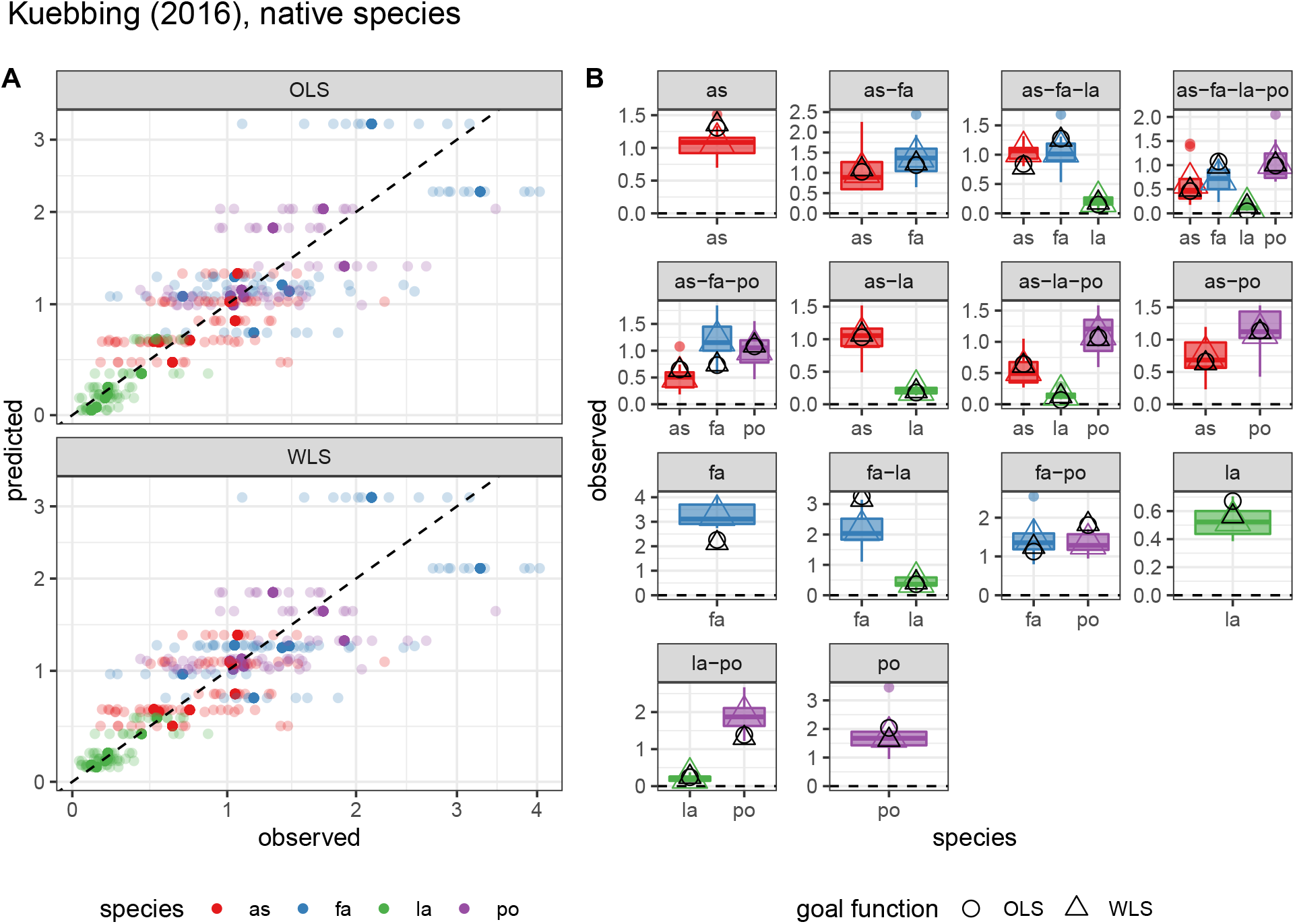
As Figure 6, but for the data of Kuebbing et al. (2015), native plants.

**Figure S16:**
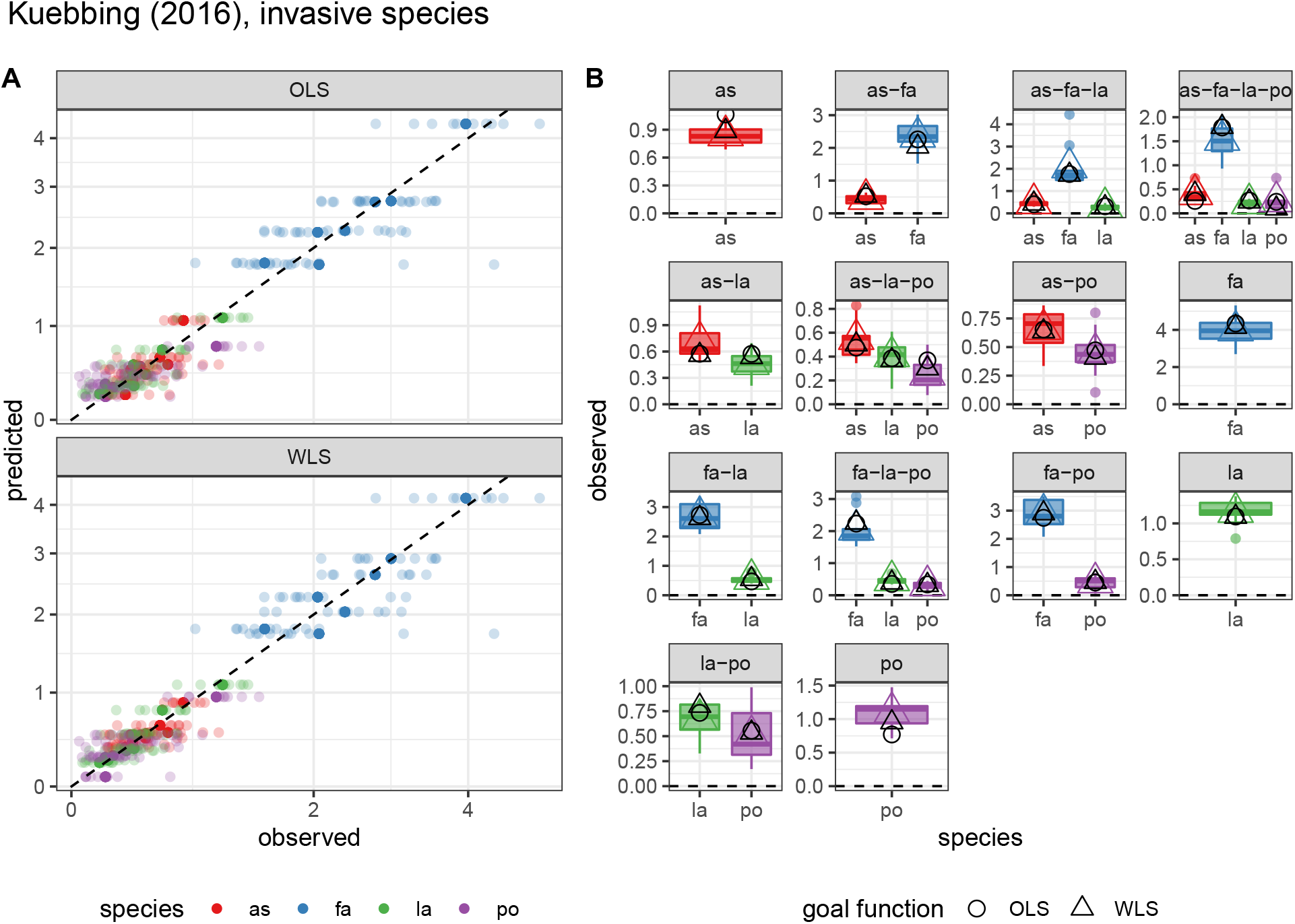
As Figure 6, but for the data of Kuebbing et al. (2015), non-native plants.

### S9. Simulations

In this section, we present results obtained when simulating noisy data and attempting to fit the “observed” data and recover the parameters used to generate the data in the first place. These simulations allow us to highlight two aspects of the approach presented here: a) WLS is superior to OLS whenever the observations on abundances display a marked mean-variance relationship; and b) while the approach presented in the main text is fundamentally based on normally-distributed deviations from the theoretical mean (as found in both OLS and WLS), one could easily extend the approach to other distributions—in which case the function to maximize would be a more generic likelihood (rather than simply minimizing the sum of squared deviations).

All the simulations presented below are based on a simple, 3-species “true” matrix of interactions:

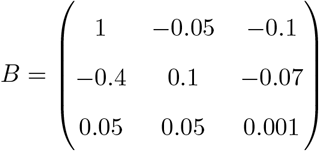

which was chosen for three reasons: a) it yields coexistence for all species in all combinations; b) it yields “true” abundances 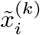 spanning a large range of values (from 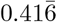 to 1000); c) it contains a mix of “mutualistic/facilitation” and “consumer-resource” relationships (signs are reversed), showing that the approach presented here extends to systems in which competition is not the only or primary mode of interaction between species.

This matrix of interactions yields a matrix of “true” values *X*, detailing the abundance each species would achieve when grown in one of the seven possible combinations of species. We then simulate the matrix of “observed” abundances 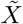 by sampling each abundance 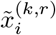 independently from a given distribution with mean 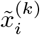. To explore distributions with prescribed mean-variance relationships, here we examine the case in which, for each community, we sample 50 noisy replicates and take 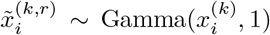, or 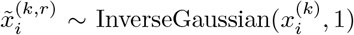. For this parameterization of the Gamma distribution, we have that the expectation 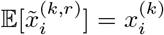, and the variance 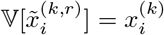, yielding a linear mean-variance relationship; for the Inverse Gaussian distribution, on the other hand, we have the same expectation 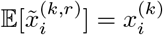, but a variance that grows rapidly with the mean, 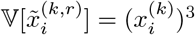.

Naturally, even if we were to choose the correct matrix *B* to fit the data, the residuals would not be normally-distributed around the corresponding means, because the shapes of these distributions depart considerably from normality (e.g., they are not symmetric around the mean). Because of this fact, and because the variance changes considerably with the mean, we expect the OLS to perform poorly—in particular, as explained in the main text, we expect low abundance values, which contribute less to the sum of squared deviations, to be fit especially poorly, while the high abundance measurements should be predicted with more accuracy. This problem should be considerably eased by performing WLS—in this case the problem of the different mean-variance relationships should be removed, while the fact that we are implicitly (wrongly) assuming normally-distributed (weighted) residuals remains. We can however attempt a different fitting routine: instead of searching for parameters minimizing the (weighted) squared deviations from the mean, we can specify an alternative form of the distribution and attempt to identify the maximum-likelihood parameters by searching for the matrix *B* maximizing the product of the densities of all observations, when either assuming that samples are taken from a Gamma distribution with shape parameter 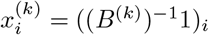 and rate parameter 1, or alternatively from an Inverse Gaussian distribution with mean parameter 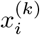 and shape parameter 1. Note that, because these are exactly the distributions we’ve used to generate the data, when given a sufficiently large number of replicates per community, our search should converge to the true matrix *B*.

The results are presented in Figs. 17-20. First, we examine the fitted abundances vs. “observed” abundances for the two distributions and three fitting routines (OLS, WLS, or likelihood-based). In Fig. S17 we show the results for simulations using a Gamma distribution: as expected, the OLS approach performs poorly, failing especially to fit the low-abundance points; the fit is greatly improved by WLS, which in fact fits the data almost as well as when maximizing the likelihood choosing the “right” distribution. The same is true for data sampled from the Inverse Gaussian distribution (Fig. S18).

Next, we consider the quality of the parameter inference: how does the inferred *B* compare to that used to generate the data? In Fig. S19 (Gamma) and S20 (Inverse Gaussian), we show that indeed the parameter inference obtained using WLS is much superior to that obtained under OLS, further demonstrating the utility of this approach.

**Figure S17:**
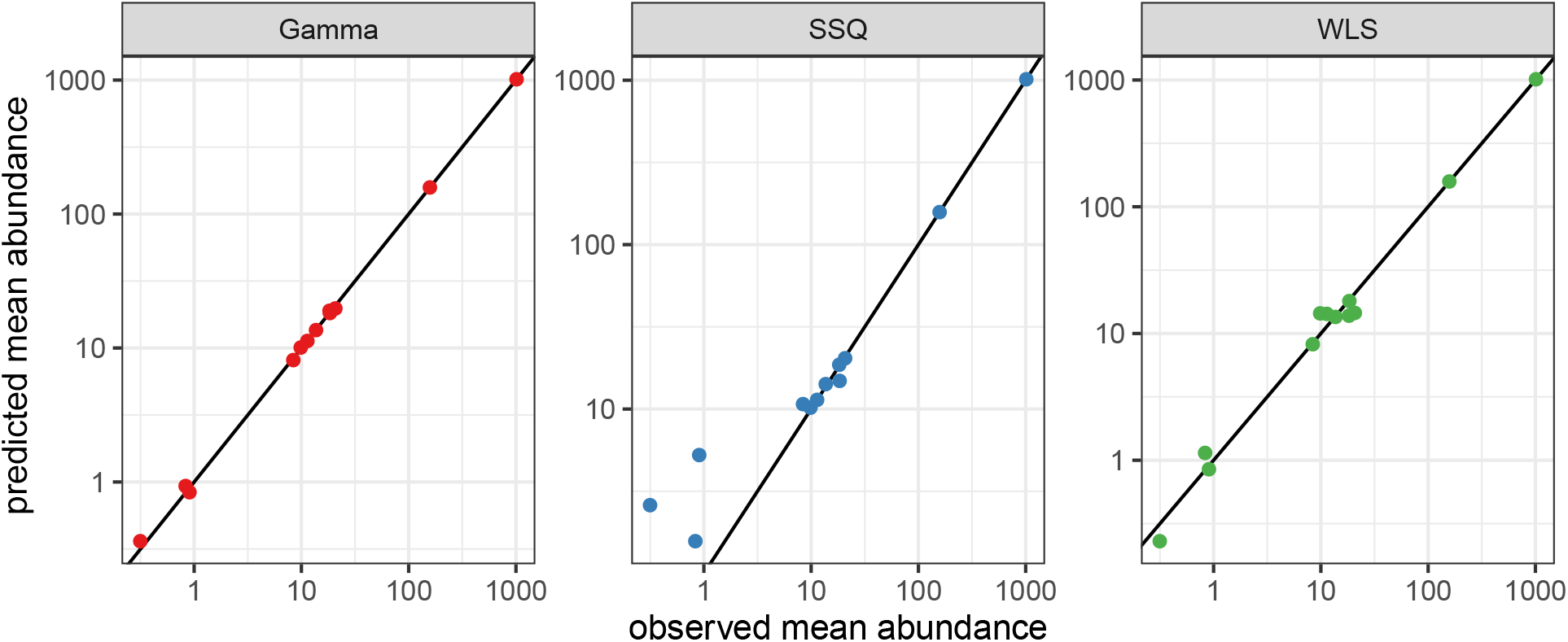
Fitted vs. observed densities for simulations in which replicates for a given system are sampled from a Gamma distribution. For each community, we plot the average abundance taken over the fifty replicates.

**Figure S18:**
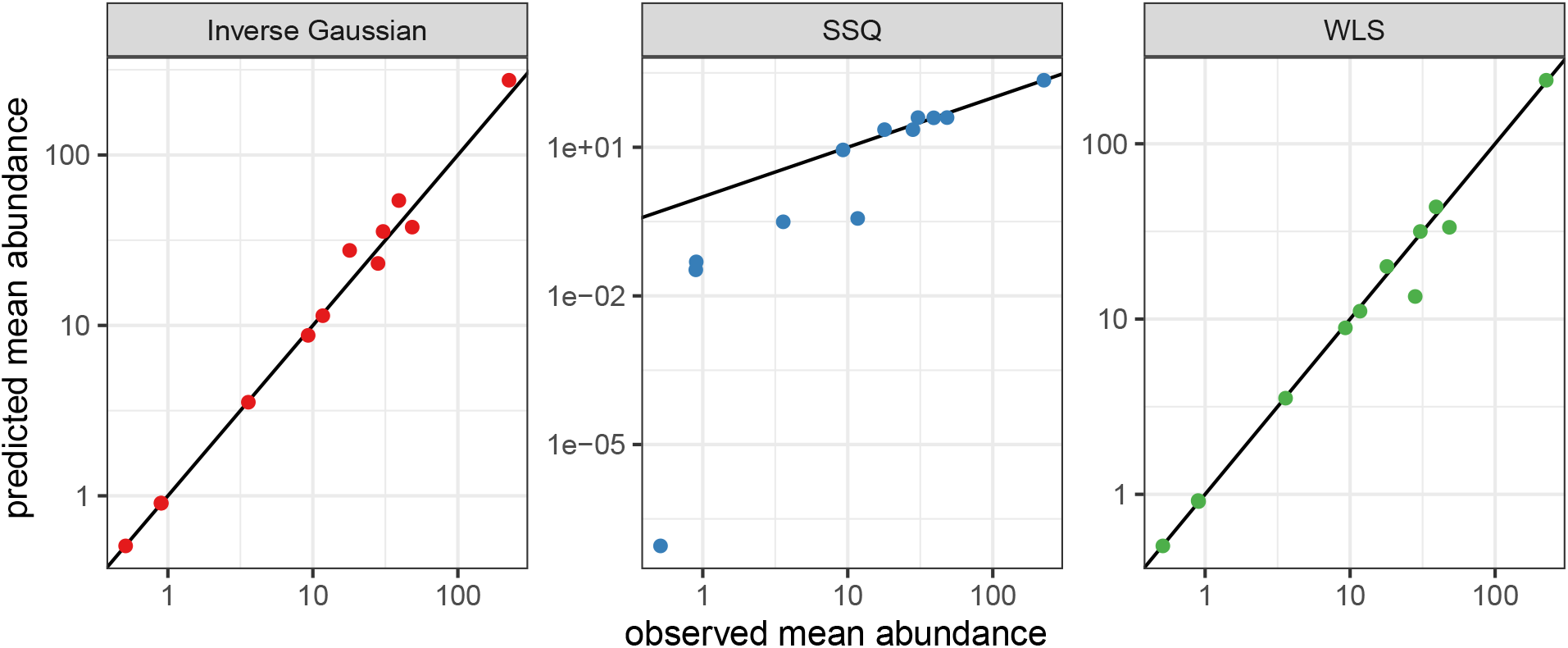
As Fig. S17, but simulating data sampled from an Inverse Gaussian distribution.

**Figure S19:**
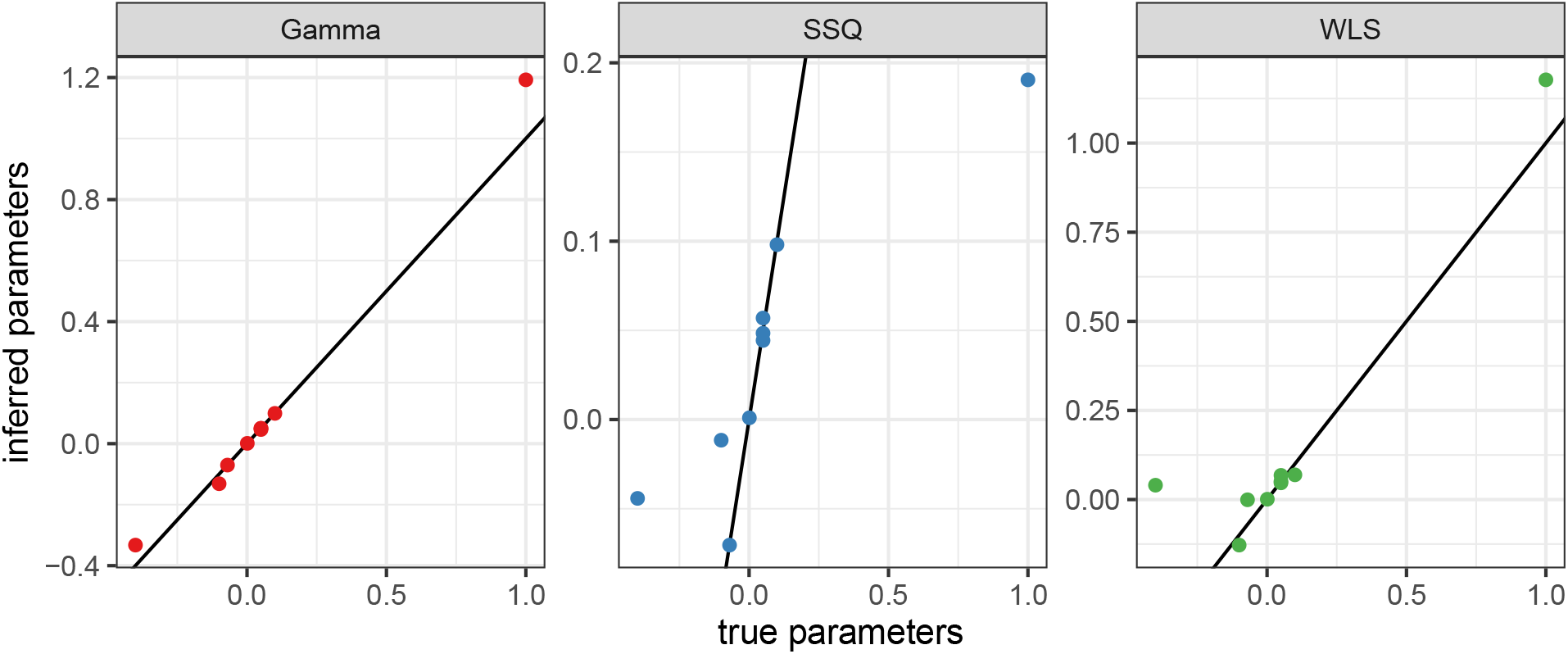
Inferred vs. true parameters for simulations in which replicates for a given system are sampled from a Gamma distribution. For each parameter, we plot the inferred value using the specified approach and fifty replicates for each community.

**Figure S20:**
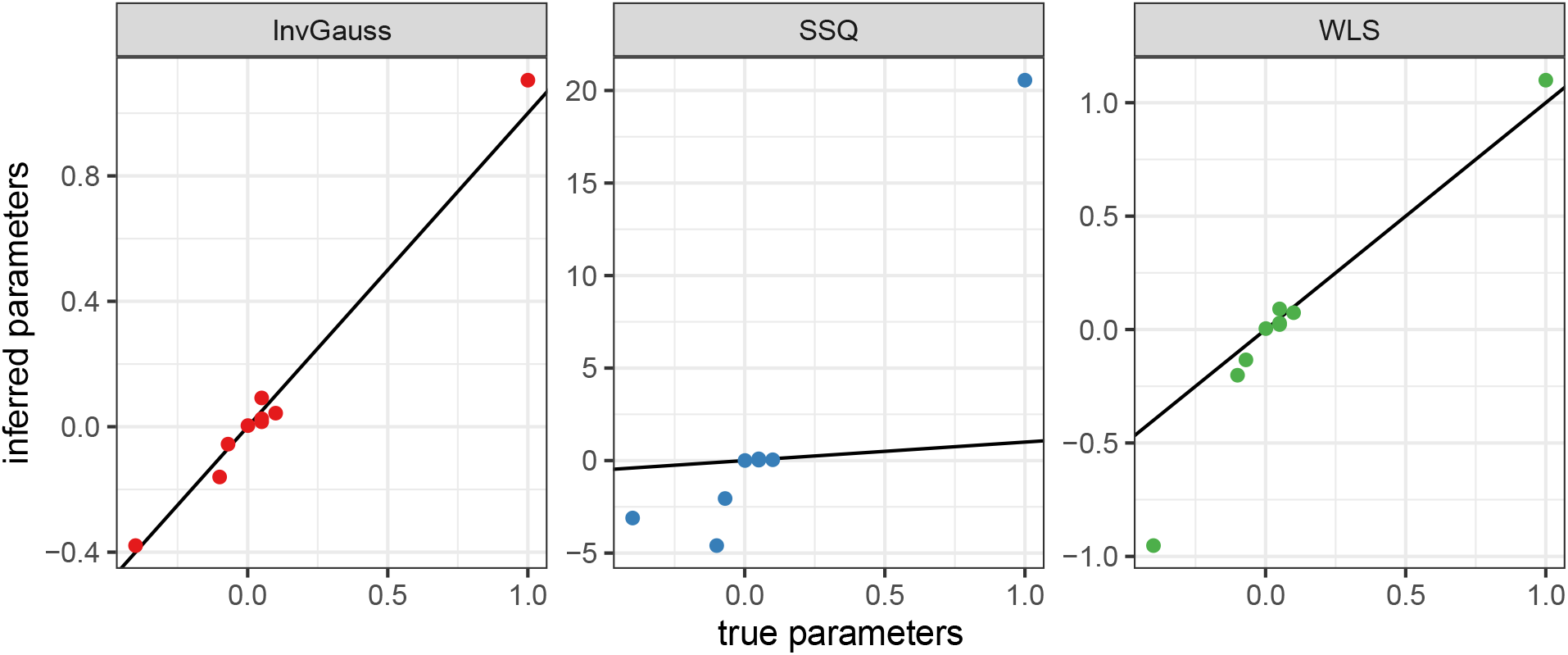
As Fig. S19, but simulating data sampled from an Inverse Gaussian distribution.

